# The importance of environmentally-acquired bacterial symbionts for the squash bug (*Anasa tristis*), a significant agricultural pest

**DOI:** 10.1101/2021.07.14.452367

**Authors:** Tarik S. Acevedo, Gregory P. Fricker, Justine R. Garcia, Tiffanie Alcaide, Aileen Berasategui, Kayla S. Stoy, Nicole M. Gerardo

## Abstract

Most insects maintain associations with microbes that shape their ecology and evolution. Such symbioses have important applied implications when the associated insects are pests or vectors of disease. The squash bug, *Anasa tristis* (Coreoidea: Coreidae), is a significant pest of human agriculture in its own right and also causes damage to crops due to its capacity to transmit a bacterial plant pathogen. Here, we demonstrate that complete understanding of these insects requires consideration of their association with bacterial symbionts in the family Burkholderiaceae. Isolation and sequencing of bacteria housed in midgut crypts in these insects indicates that these bacteria are consistent and dominant members of the crypt-associated bacterial communities. These symbionts are closely related to *Caballeronia* spp. associated other true bugs in the superfamiles Lygaeoidea and Coreoidea. Fitness assays with representative Burkholderiaceae strains indicate that the association can significantly increase survival and decrease development time, though strains do vary in the benefits that they confer to their hosts, with *Caballeronia* spp. providing the greatest benefit. Experiments designed to assess transmission mode indicate that unlike many other beneficial insect symbionts, the bacteria are not acquired from parents before or after hatching but are instead acquired from the environment after molting to a later development stage. The bacteria do, however, have the capacity to escape adults to be transmitted to later generations, leaving the possibility for a combination of indirect vertical and horizontal transmission.

## 1 Introduction

Microbial symbionts can increase their hosts’ fitness by provisioning them with nutrients or by protecting them against pathogens, parasites, and predators (Gerardo and Parker, 2014; Haine, 2008; Moran, 2006). Association with microbial symbionts, therefore, significantly alters the ecology of most hosts, which can have important applied implications when those hosts are significant pests or are vectors of disease. Indeed, some suggested methods for future control of pest and disease vectors rely on alteration of the host-symbiont association (Chuche et al., 2016; Douglas, 2007; Mendiola et al., 2020).

The squash bug, *Anasa tristis* De Greer (Heteroptera: Coreidae; Fig. 1a), is a devastating plant pest of the Cucurbitaceae family (Beard, 1940; Cook and Neal, 1999), preferentially feeding on squash and pumpkins (Bonjour and Fargo, 1989; Nechols, 1987). As sap-feeders, the bugs pierce plant tissue and cause damage to xylem transport (Neal, 1993). Furthermore, *A. tristis* is a natural reservoir and vector of the bacterium *Serratia marcescens*, the causal agent of Cucurbit Yellow Vine Disease (CYVD) (Avila et al., 1998; Bruton et al., 1998, 2003). Like many other species in the order Heteroptera, *A. tristis* possesses terminal midgut structures (Fig. 1b, c), known as crypts or ceca. The crypts house bacterial symbionts in other heteropteran species (Boucias et al., 2012; Fukatsu and Hosokawa, 2002; Garcia et al., 2014; Itoh et al., 2014; Kikuchi et al., 2005, 2011; Ohbayashi et al., 2015; Olivier-Espejel et al., 2011; Takeshita et al., 2015). In many families within the Coreoidea and Lygaoidea superfamiles and in the Largidae family within the Pyrrhocoroidea superfamily, the predominant symbionts within the crypts are bacteria from the Burkholderiaceae family (Boucias et al., 2012; Garcia et al., 2014; Gordon et al., 2016; Itoh et al., 2014; Kikuchi et al., 2005, 2011a; Ohbayashi et al., 2019b; Olivier-Espejel et al., 2011; Ravenscraft et al., 2020; Sudakaran et al., 2015; Takeshita et al., 2015; Xu et al., 2016a). These symbionts most commonly have been referred to as *Burkholderia*, though evidence supports reclassification of these symbionts into the genus *Caballeronia* (Beukes et al., 2017; Dobritsa and Samadpour, 2016, 2019; Takeshita and Kikuchi, 2020), and hereafter we will refer to them as *Caballeronia* spp.. Other bacteria can inhabit crypts as well. In some species, these bacteria are co-inhabitants with *Caballeronia* (Boucias et al., 2012; Fukatsu and Hosokawa, 2002; Garcia et al., 2014; Gordon et al., 2016), while in others they are alternatives to a *Caballeronia* association (Nishino et al., 2021; Takeshita and Kikuchi, 2020).

**Figure 1.**
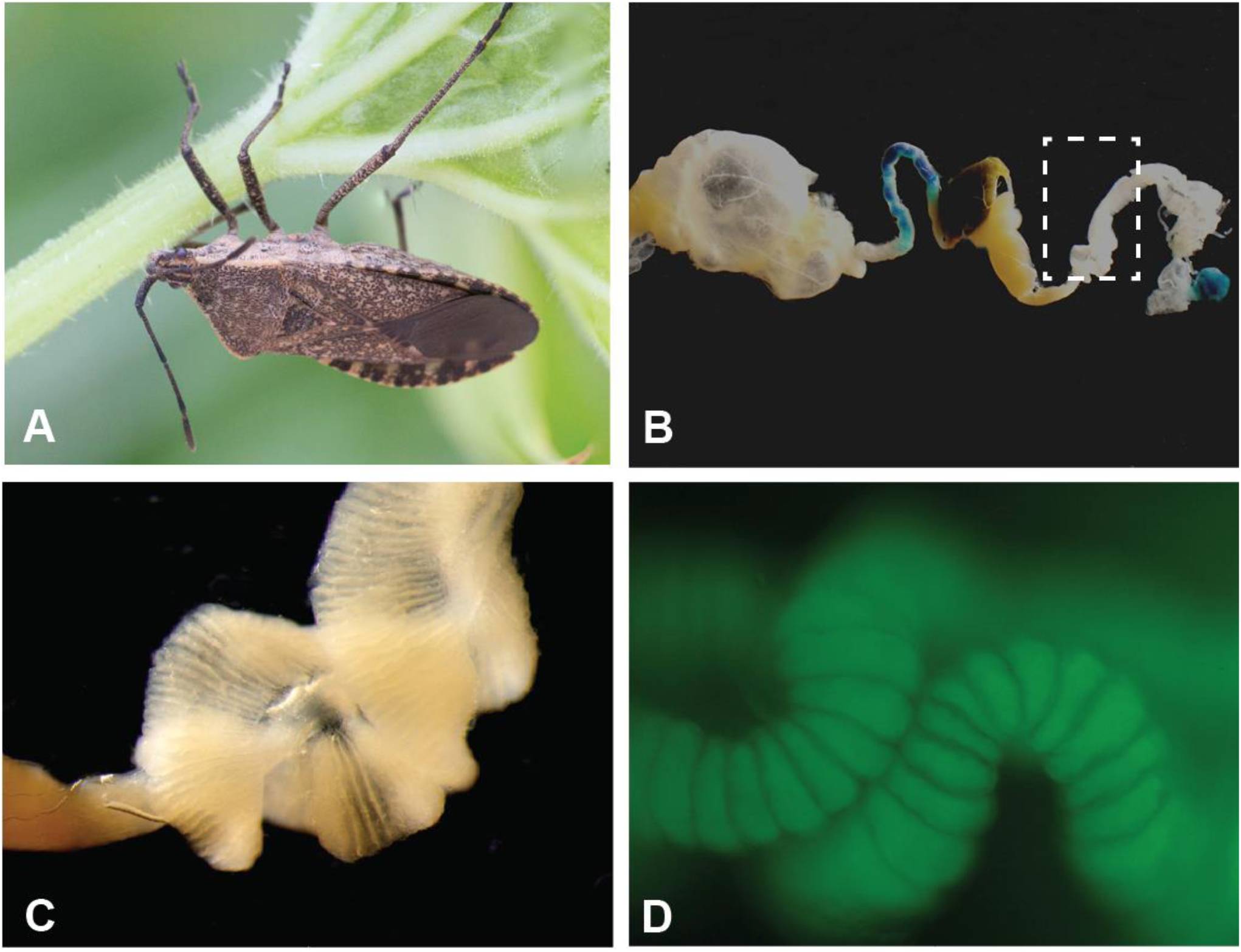
Specialized midgut crypts of the agricultural pest *Anasa tristis*. (A) Adult *A. tristis*. (B) Entire midgut of adult *A. tristis*, with box highlighting M4 section. (C) Close-up of M4 section, which has the highly invaginated surface typical of midgut symbiont crypts in other true bug species. (D) Close up of crypts with GFP-labeled *Caballeronia* sp. bacteria.

*Caballeronia* symbionts can provide their hosts with several benefits. These include faster development, increased survival and reproduction, increased body size, stronger immune-based defense and insecticide resistance (Boucias et al., 2012; Garcia et al., 2014; Itoh et al., 2018; Kikuchi et al., 2007, 2012; Kikuchi and Fukatsu, 2014; Kim et al., 2015; Lee et al., 2017; Olivier-Espejel et al., 2011). The benefits conferred differ based on the insect species and, in at least one species, based on the bacterial strain (Itoh et al., 2019; Kikuchi et al., 2012; Ravenscraft et al., 2020).

It is well documented that symbionts can alter the life history and food preference of crop pests (Hosokawa et al., 2007; Toju and Fukatsu, 2010), as well as alter transmission of pathogens by insect vectors (Arora and Douglas, 2017; Crotti et al., 2012; Su et al., 2013; Weiss and Aksoy, 2011). It is currently unknown whether *A. tristis* harbors beneficial bacterial symbionts. Here, we identify the bacteria within the midgut crypts of *A. tristis* through culture-independent and culture-dependent 16S rRNA sequencing. We then determine how these bacteria are transmitted between squash bug generations and how they affect host fitness. We demonstrate that crypt communities are dominated by *Caballeronia* spp. that are predominantly acquired from the environment each generation, leaving the possibility for individual insects to pick up strains with alternative fitness traits. Controlled infection experiments demonstrate that association with these bacteria can significantly increase host survival and decrease development time, and that bacterial strains vary in their ability to colonize and provide benefits to their hosts. Taken together, these data suggest that these bacteria are a primary, beneficial symbiont of these agricultural pests and vectors of plant disease.

## 2 Materials and Methods

### 2.1 Characterization of Crypt Bacterial Community through Culture-Independent Sequencing

*A. tristis* adults were collected from Crystal Organic Farm in Georgia, United States from crookneck squash plants (*Cucurbita pepo*) in the summer of 2014. Insects were transported to the laboratory and either dissected immediately or housed with plant material and conspecifics from their collection site until dissection. Squash bugs were anesthetized with CO_2_ gas, sacrificed, surface-sterilized in 95% ethanol for five minutes and rinsed in sterile Carlson’s solution (Mitsuhashi, 2002). We dissected the midgut from 10 adults: the m1 and m4 organs from the midgut were collected from five adults, and the entire midgut was dissected from the remaining five adults. These tissues were rinsed with Carlson’s solution and then crushed with a micro-pestle in Carlson’s solution. DNA was extracted from each sample using a cetyl trimethylammonium bromide (CTAB) protocol. Specifically, an equal volume of CTAB was added to each sample and then incubated at 60° C for one hour. Sodium dodecylsulfate (SDS) was added to a final concentration of 2%, and samples were incubated at 60° C for one hour. Nucleic acids were extracted with an equal volume of 24:24:1 phenol:chloroform:isoamyl alcohol and then extracted twice with an equal volume of chloroform. Two volumes of cold 99.5% ethanol and 0.1 volume of NaOAc were added, and then samples were placed at −20° C to precipitate overnight. Pelleted DNA was washed with 75% ethanol, dried, and resuspended in molecular grade water.

We surveyed the bacterial microbiome by amplifying the V4 region of the 16S rRNA gene using dual-indexed primers (Kozich et al., 2013). Each primer consisted of Illumina adapter, index, pad, link, and V4-specific sequences. Specifically, the V4 16S forward primer (5’-GTGCCAGCMGCCGCGGTAA-3’) was combined with i5 index sequences and the V4 16S reverse primer (5’-GGACTACHVGGGTWTCTAAT-3’) was combined with i7 index sequences. These primers resulted in a 250 bp amplicon. The 5 PRIME Master*Taq* PCR kit was used to amplify the V4 region of the 16S rRNA gene in 25 uL reactions containing (final concentrations) 0.5X 5 PRIME Master*Taq* Buffer with Mg^2+^, 200 μM dNTPs, 400 nM each forward and reverse primers, 1X *Taq*master PCR enhancer, 1 mM MgCl_2_, 1 U *Taq* polymerase and genomic DNA. Reactions were denatured at 94° C for 2 mins followed by 30 cycles of: 1) 94° C for 20 s, 2) either 55° C or 60° C for 15 s, 3) and 72° C for 5 min, with a final elongation step at 72° C for 10 min. Amplicons were purified with the Qiagen PCR Purification kit, eluted with molecular grade water, and quantified with the Qubit dsDNA BR Assay kit using the Qubit Fluorometer. Samples were pooled in equimolar concentrations and then run on an Illumina MiSeq with the MiSeq Reagent Kit v2.

MiSeq data were analyzed following a standard pipeline (Kozich et al., 2013) in mothur (Schloss et al., 2009). Paired-end reads were assembled into contigs and screened to remove contigs that were the wrong length, had ambiguous reads or had more than eight homopolymers. The remaining contigs were dereplicated, aligned to sequences from the nonredundant SILVA database (release 119), and pre-clustered into OTUs using a 97% similarity threshold. Chimeras (identified by VSEARCH) were removed. Sequences were taxonomically classified using the Wang method with template sequences from the Ribosomal Database Project (training set 18 released June 2020). Sequences identified as archaea, eukaryote, mitochondria, or chloroplast were removed, and the remaining sequences were clustered into phylotypes based on their genus-level classification. Phyloseq was used to visualize MiSeq data, and vegan was used to perform Permutational MANOVA (PERMANOVA) statistical tests using Jaccard distance with 10,000 permutations.

### 2.2 Bacterial Isolation and Identification via Culture-Dependent Sequencing

*A. tristis* nymphs and adults were collected from plots containing squash and zucchini plants in six states within the United States (Arizona, Indiana, Florida, Georgia, Missouri and North Carolina). Insects were transported to the laboratory and either immediately dissected or housed with plant material and conspecifics from their collection site until dissection. We dissected midgut crypts (the M4 section of the *A. tristis* midgut; Fig. 1b, c) from 55 adults, one fifth instar, two fourth instars, and two third instars. We then crushed the crypts in either Carlson’s solution or 1X phosphate buffered saline (PBS). One whole second instar nymph was crushed in Carlson’s solution, as the midgut was too underdeveloped to dissect. Bacteria were cultivated on Luria-Bertani (LB) agar (incubated for 48 hours at 27° C), a medium on which symbionts from other coreid bugs readily grow (Garcia et al., 2014; Kikuchi et al., 2007). Isolated bacteria were stored as glycerol stocks at −80° C.

We used sequencing to identify one to twelve bacterial isolates from each of the 61 bugs sampled. We extracted DNA from bacterial isolates collected prior to 2019 from Georgia and Missouri using a CTAB extraction protocol. Briefly, overnight cultures of each bacterial isolate were lysed by incubating them with 2% CTAB buffer and 10% sodium dodecylsulfate (SDS), and then nucleic acids were extracted once with phenol:chloroform:isoamyl alcohol (25:24:1) and twice with chloroform. DNA was recovered by ethanol precipitation at −20° C We extracted DNA from all other bacterial isolates by boiling a single colony of each isolate in molecular-grade water (ten minutes at 95° C).We amplified a portion of the 16S rRNA gene from each isolate using the MasterTaq® kit (5 PRIME) and universal bacteria primers 27F (5’ AGA GTT TGA TCC TGG CTC AG 3’) and 1492R (5’ GGT TAC CTT GTT ACG ACT T 3’) (Lane, 1991). PCR amplifications were performed with an initial 4 min denaturing at 94° C followed by 36 cycles of denaturing for 30 s at 94° C, annealing for 30 s at 55° C and extending for 1 min at 72° C, with a final 1 min extension at 72° C. Amplicons were purified using the QIAquick® PCR Purification Kit and sequenced with the forward primer. All sequences are deposited in Genbank (accession numbers KT259132 - KT259191, KX239751 – KX239768, MH636869 – MH636872 and MZ264232-MZ264276) We trimmed the sequences from prior to 2019 using CodonCode Aligner 5.1.5 and all other sequences using SeqMan Pro (DNA STAR Navigator v. 16). Trimmed sequences greater than 500 bps in length were identified to genus through comparison to RDP, the Ribosomal Database Project (training set 18 released June 2020). Information is presented for sequences that had a greater than 60% match at the genus level. *Anasa*-derived Burkholderiaceae sequences with a length greater than 600 bps, along with additional Burkholderiaceae sequences (see Supplemental Fig. 1 for details) were aligned and curated with MAFFT (v 7.407_1) and BMGE (v 1.12.1) as implemented at NGphylogeny.fr (Lemoine et al., 2019). Through NGphylogeny.fr, we used SMS (v 1.8.1) to estimate the best model of evolution and PhyML (v 3.3.1) to construct a maximum likelihood phylogeny with aLRT-SH branch support (Ehman et al., 2018; Guindon et al., 2010; Lefort et al., 2017). The tree was visualized using Figtree v1.4.2.

### 2.3 Symbiont Transmission

To determine symbiont transmission mode, and for all other experiments, we established colonies of *Anasa tristis* in the laboratory. These colonies were originally started by collecting individuals of all instars from gardens and farms in Georgia. Insects are reared in net tents containing multiple pairs of adults and one to two potted squash plants. Plants are changed regularly, and occasionally the colonies are transferred to new tents. Eggs are collected regularly for experiments.

We surfaced sterilized eggs with a 70% ethanol wash for two minutes followed by a 10% bleach wash for two minutes and a ten second rinse with sterile water. Emerging first instar nymphs were fed on a piece of surface sterilized acorn squash in a bleach-cleaned plastic box until they molted into the second instar. We then starved the second instars nymphs for 12 hours and gave them access to a parafilm-covered Petri dish (< 20 nymphs per dish) with sterile water, 1% blue food-grade dye, which allowed us to assess if the individuals had fed on the solution, and Green Fluorescent Protein (GFP)-labeled *Caballeronia* sp. SQ4a (Fig. 3) at a concentration of ~10^7^ cells/mL. *Caballeronia* sp. SQ4a was isolated from an *A. tristis* adult collected at Oakhurst Garden and fluorescently labeled with GFP using a previously described triparental mating protocol (Kikuchi and Fukatsu, 2014). After 24 hours, we removed the Petri dish and replaced it with a surface sterilized acorn squash piece for 24 hours. To minimize potential exposure to other environmental bacteria, nymphs were then placed into semi-sterile mesh tents (12 in × 12 in × 12 in by Raising Butterflies) with axenically grown squash plants. We prepared these semi-sterile environments by: 1) sterilizing the tents and water reservoirs in an autoclave; 2) washing and vortexing each squash seed in 100% bleach for five minutes; 3) germinating the seeds in sterilized boxes; and, 4) transferring the seeds to autoclaved pots with sterile perlite, a soil substitute. These pots extended into reservoirs filled with sterile water with Botanicare Pure Blend Pro Grow Organic Fertilizer (Fig. S2). The nymphs were reared in these environments until adulthood.

We then placed these *A. tristis* adults that had been fed GFP-labeled *Caballeronia* sp. SQ4a as nymphs into two mesh tents, similar to those described above but with one or two squash plants grown in non-sterile potting soil. The tents contained three to four adults each (two female and two male adults in tent 1; one female and two male adults in tent 2). These adults mated and laid eggs freely. Eggs and nymphs of all instar stages were periodically removed from both tents to test for the presence or absence of GFP-labeled bacteria. The only mechanism by which these bugs could acquire GFP-labeled bacteria is if it was passed directly from a parent or if it escaped a bacteria-inoculated adult and then was acquired from the environment. Since the soil and plants were not sterile, individuals had the opportunity to pick up alternative strains of non-GFP-labeled bacteria as well.

We screened for the presence of GFP-labeled bacteria in eggs and nymphs by visualizing GFP-labeled bacterial colonies isolated from eggs and insects reared in the tents. We surface sterilized some eggs and all nymphs by washing each in 99.5% ethanol for 30 seconds; both non-sterile and surface sterilized eggs were utilized to determine whether bacteria came from within the egg or from the egg surface. Eggs (two per sample) and first instar nymphs were crushed with a pestle in 200 μL of Carlson’s solution. Second, third, and fourth instar nymphs were crushed in 300 μL of Carlson’s solution. We plated 50 - 100 μL of each solution, in triplicate, onto LB agar with 30 μg/mL kanamycin and incubated the plates at 27° C for two days. We then assessed presence of GFP-labeled colonies using a fluorescent microscope.

To determine whether low numbers of GFP-labeled bacteria could be detected using the above plating method, we washed eggs in bacterial solutions of known concentrations of GFP-labeled *Caballeronia* and quantified the bacteria on these eggs using the above methods. For each concentration tested, six surface-sterilized eggs were put into a 1.5 mL tube. We then added 800 μL of a solution with a concentration of approximately 5 × 10^3^, 5 × 10^2^, 5 × 10^1^, or 5 CFUs of SQ4a per μL (concentrations determined by plating three replicates of 2 μL of the inoculation solution). After 10 minutes, eggs were removed from the solution and allowed to air dry. We plated three replicates per sample. We detected bacteria in all samples washed in solutions of more than 5 CFUs per μL (Supplemental Table S3).

### 2.4 Symbiont Uptake and Colonization

In the above transmission experiment, we did not find GFP-labeled *Caballeronia* associated with any egg or first instar samples. To determine whether first instars were capable of establishing a *Caballeronia* infection upon ingestion, we reared first instars from surface sterilized eggs and then exposed them to GFP-labeled *Caballeronia* sp. SQ4a using several methods. The first method consisted of placing nymphs on a sterile dental cotton roll extending out of a small Petri dish. The dish was filled with bacterial solution and capped with parafilm. Nymphs were allowed to walk on and probe the resulting saturated cotton for ten minutes. The second method consisted of nymphs placed in a small Petri dish with five 2 μL droplets of bacterial solution. Only nymphs that were seen probing a droplet were monitored for subsequent symbiont establishment. Similar to the first method, the third method allowed nymphs to walk on bacteria-saturated cotton rolls for ten minutes, but they were prodded with a sterile inoculation loop to prevent them from probing the cotton; this method was used to determine if the bug would establish the symbiosis once the bacteria were on the surface of their exoskeleton. As a comparison, we used similar procedures to inoculate second instar nymphs using the first two methods. In total, we exposed 18 first instars and 15 second instars using the ‘cotton’ method, seven first instars and five second instars using the ‘droplet’ method, and seven first instars using the ‘surface’ method. Nymphs were sacrificed after molting to the next instar in order to assay whether they contained any GFP-labeled bacteria using plating methods described above.

To estimate the *Caballeronia* sp. SQ4a load carried in adults upon establishment, second instar nymphs were reared from surface sterilized eggs and fed GFP-labeled *Caballeronia* sp. SQ4a using the cotton method detailed above. Individuals were then placed on semi-sterile hydroponic plants, as described above, and reared to adulthood. Once they reached adulthood, we dissected midgut crypts from 10 females and 13 males, and, as described above, dilution plated these samples onto Yeast extract and Glucose (YG) media to estimate the total CFUs per crypt that were GFP-labeled.

### 2.5 Fitness Benefits of Symbiosis with a Focal Symbiont for Host when Reared on Plants

We fed second instar nymphs on a sterile Petri dish with GFP-labeled *Caballeronia* sp. SQ4a or without GFP-labeled bacteria (water control), using protocols described above. We then replaced the diet with a piece of sterile acorn squash, and fed the inoculated insects on fruit for 1-10 days (mean 4.6 days for bacteria treatment, 4.5 days for control treatment) before moving them to plants. We set up five replicate plant tents, as described above, with *Caballeronia* sp. SQ4a inoculated second instars (n = 8, 10, 11, 16, 17 individuals per tent) and six replicate plant tents with water inoculated (control) second instars (n = 6, 8, 9, 10, 14, 16 individuals per tent). Insects were reared in a climate-controlled chamber at 28° C (16 hr L:8 hr D).

Each day, we recorded the number of individuals alive and the developmental stage of live individuals in each tent. We analyzed survival over time as a step function using Kaplan-Meier survival analysis with the ‘survival’ package in R (Therneau, 2013), and analyzed differences in survival between treatments with Cox regression analysis (Klein and Moeschberger, 2003). We censored the data at the time individuals reached adulthood or, in a few cases, when individuals could not be found. We sacrificed adults and confirmed that they either contained (*Caballeronia* sp. SQ4a treatment) or did not contain (water control treatment) GFP-labeled bacteria in the crypts, using protocols described above. All *Caballeronia*-fed individuals contained GFP-labeled bacteria; no control individuals did.

By monitoring the tents daily, we determined the proportion of insects in each cage that survived and molted to third, fourth and fifth instars and to adult. These four dependent variables were analyzed separately using quasibinomial distributed Generalized Linearized Models (GLMs), with treatment (*Caballeronia* sp. SQ4a or water-fed controls) as the explanatory factor.

For each individual that survived to a given stage, we estimated development time from: hatch to third instar, hatch to fourth instar, hatch to fifth instar, and hatch to adult emergence. While one would ideally compare the time between each consecutive life stage rather than from hatch to each stage, such data were not estimable based on our counting methodology, and we recognize that these variables are overlapping and therefore correlated. These four dependent variables were analyzed separately using Wilcoxon rank sum tests, with treatment as the explanatory factor. We conducted all statistical analyses in R v3.0.2 (R Core Team, 2013).

### 2.6 Fitness Benefits of Symbiosis with a Focal Symbiont for Host when Reared on Plants versus Fruits

In the field, squash bugs feed on both plants and fruits. To determine whether host diet could influence the benefits of symbiosis, we reared sterile first instar nymphs by surface sterilizing eggs as described above. We fed emerging first instar nymphs on surface sterilized zucchini squash until they molted into the second instar. Upon molting, nymphs were randomly distributed into two treatments: (i) *Caballeronia*-infected or (ii) uninfected control. Bacterial infection was carried out with unlabeled *Caballeronia* sp. SQ4A, following the “cotton” method described above. Non-infected nymphs were fed water as a control. Following the infection treatment, nymphs were further divided into two new groups representing two different feeding substrates, plant and fruit. This resulted in an experimental design with four treatments: (i) *Caballeronia*-infected nymphs on plants (n = 41), (ii) *Caballeronia*-infected nymphs on fruit (n = 38), (iii) non-infected nymphs on plants (n = 41) and (iv) non-infected nymphs on fruit (n = 38). Nymphs in the plant treatment were reared in groups of 8 to 15 on squash plants growing in nutrient water within mesh tents as described above. Nymphs in the fruit treatment were placed in groups of not more than ten individuals in plastic boxes containing a piece of surface-sterilized acorn squash. Each treatment was conducted in quadruplicate. The experiment ran for 29 days in a climate-controlled chamber at 28° C (16 hr L:8 hr D).

Fitness was assessed by monitoring the tents and boxes daily and measuring the same fitness parameters described above. Statistical analyses were carried out as described above for analysis of survival and proportion to each instar data, with infection treatment (*Caballeronia* sp. SQ4a or water-fed controls) and food substrate (plant or fruit) as explanatory factors. Development time from hatch to each instar was analyzed using quasipoisson distributed Generalized Linearized Models (GLMs); as above, this was done separately for the time to third instar, fourth instar, fifth instar and adult. Tukey’s post-hoc tests were used to calculate differences between the groups using the R package “multcomp” (Bretz et al., 2010).

### 2.7 Variation in Fitness Benefits Conferred by Alternative Bacteria

Second instar nymphs (n = 60 to 76 per treatment) were fed one of four different GFP-labeled bacterial strains (SQ4a, A33_M4_a, SMT4a, BHJ32i) or a water control, using methods describe above. A33_M4_a is a *Caballeronia* sp. isolated from the crypt of an adult *A. tristis* from Crystal Organic Farm (Newborn, Georgia, United States), and SMT4a is a *Paraburkholderia* sp. (Burkholderiaceae), isolated from soil at a site with other *Caballeronia*-associated true bug species in the genus *Alydus* (Garcia et al., 2014); both strains are indicated in Fig. 3. BHJ32i, a *Cupriavidus* sp. (Burkholderiaceae), was isolated from the crypt region of an *Alydus tomentosus* adult (Garcia et al., 2014). Hereafter, for brevity, treatments are referred to as “SQ4a”, “A33”, “SMT”, “BHJ” and “H2O.” After a 24hr rest period following bacterial feeding, nymphs were placed in sterilized boxes (five nymphs per box) with surface sterilized zucchini squash. Development was recorded daily, and the fruit was replaced daily. Upon reaching the adult stage, recently molted adults were weighed and photographed. From photographs, we estimated four measures of body size (scutellum width, pronotum width, posterior tibia length and body length) using ImageJ. Live adults were then isolated from one another. Adults that died before being dissected were also weighed and imaged for size measurements within 24 hr of dying. To quantify bacterial load, adults were dissected as previously described 2-5 days after molting, and their crypts were crushed with a pestle in 500 uL Carlson’s solution. Serial dilution was performed in LB broth, and cells were plated onto YG plates. Colony forming units in the crypts were estimated from two replicate plates at the most appropriate serial dilution. Statistical analyses of survival and development data were carried out as described above. Log-transformed pronotal width, square root-transformed adult weight data and log-transformed colony forming unit data were analyzed with ANOVAs, with sex and bacterial inoculation treatment as factors. Tukey’s multiple comparisons were used to further assess the impacts of bacterial treatment on pronotal width and on bacterial colonization.

## 3 Results

### 3.1 Characterization of Crypt Bacterial Community through Culture-Independent Sequencing

*Caballeronia* was the main constituent of the midgut crypts (m4 organ) in four of the five adult squash bugs we sampled (Fig. 2A) and accounted for an average of 68.0% of the MiSeq reads recovered from the crypts. The crypts of one insect, SBA.36, were instead dominated by unclassified Enterobacterales (80.4%), but unclassified Burkholderiaceae was the second most abundant phylotype (17.5%) in this sample. *Caballeronia* was also present in the m1organs and whole midguts, but at lower relative levels than in the crypts (Fig. 2B). The microbiomes of the m1 and m4 organs were significantly different (PERMANOVA, F = 2.65, d.f. = 1, *P* = 0.039).

**Figure 2.**
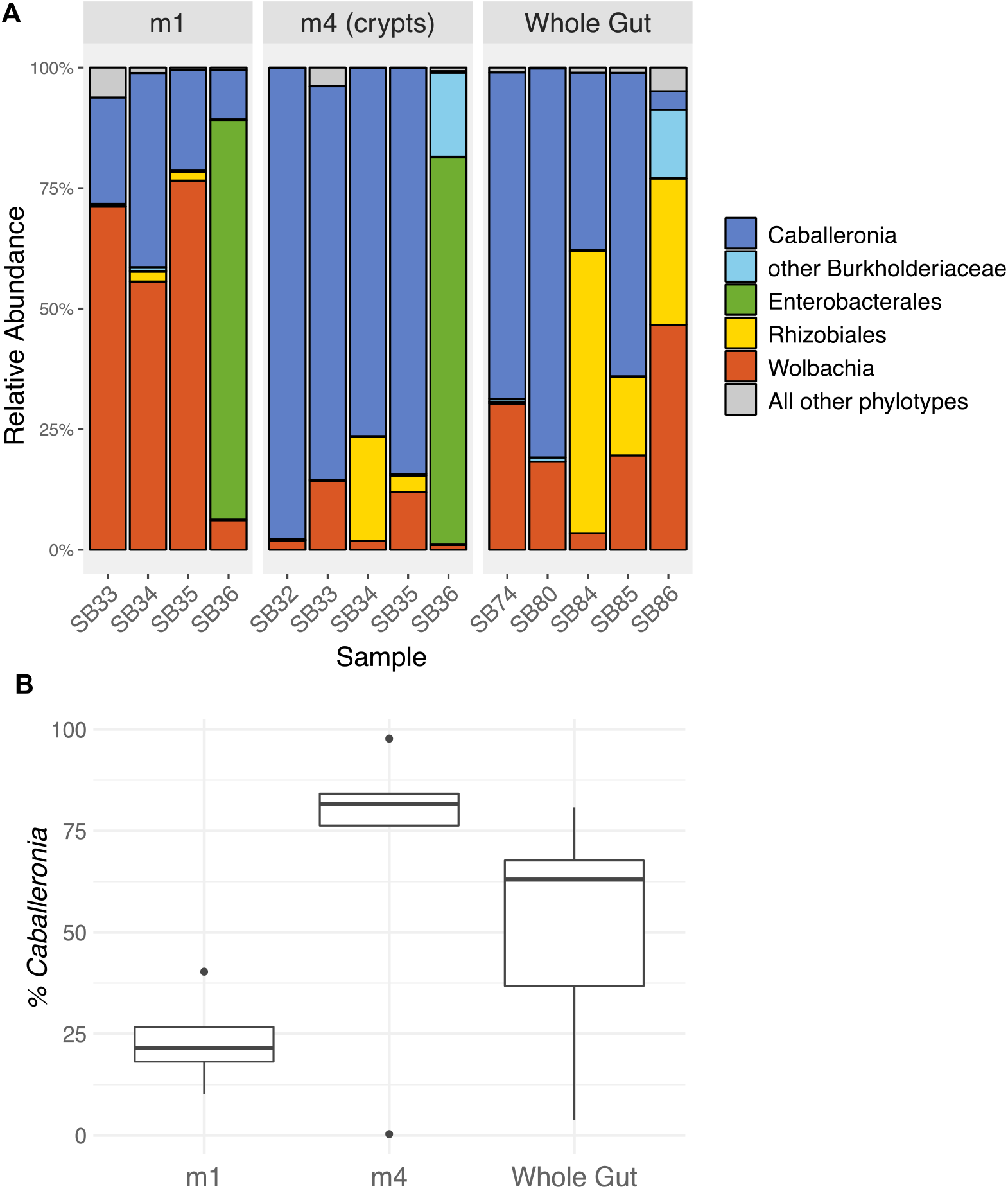
Relative abundance of 16S rRNA gene MiSeq reads from adult squash bug midgut sections. **(A)** MiSeq reads are grouped into genus-level phylotypes and labeled with the lowest certain taxonomic classification according to RDP classification. The five most abundant phylotypes are shown and all other phylotypes are combined. The m1 and m4 organ samples were taken from the same individuals. The m1 organ for sample SB32 is not included because it failed to amplify. **(B)** Percentage of reads classified as *Caballeronia* in each midgut sample. Boxplots indicate the interquartile range with a box, the median as a bold line within the box, and outliers as points. The whiskers indicate the minimum and maximum values up to 1.5x the interquartile range. Values beyond 1.5x the interquartile range are indicated as outliers.

### 3.2 Bacterial Isolation and Identification via Culture-Dependent Sequencing

We isolated and successfully sequenced bacteria from 61 individuals. Of the 128 bacteria sequenced and identified to genus, 114 were identified as *Caballeronia* (Fig. 1; Supplemental Table 1). The remaining sequences were identified as *Acinetobacter, Bacillus*, *Enterococcus*, *Klebsiella*, *Paenibacillus*, *Pseudomonas, Serratia, Staphylococcus* and *Stenotrophomonas* (Supplemental Table 2). Most of these non-*Caballeronia* isolates were recovered from M4 crypt samples, though the *Serratia* isolates were from a whole body, second instar nymph sample and thus may or may not have been living within the crypts. Both *Caballeronia* and non-*Caballeronia* species were sequenced from the midgut crypts of five individuals.

We estimated a phylogeny with the *Caballeronia* 16S rRNA sequences from *A. tristis*, as well as 16S rRNA sequences of representatives of the genera *Paraburkholderia* and *Burkholderia*. All 114 *A. tristis Caballeronia* sequences grouped with *Caballeronia* isolated from the midguts of other bug species and from plants (Fig. 3, Supplemental Fig. 1). Detectable *Caballeronia* co-infection with strains from multiple clades was rare within sampled *A. tristis*; if we consider the *A. tristis* symbionts to separate into four clades (Fig. 3), only 5 of 59 *A. tristis* were detected to have co-infections with *Caballeronia* from two or more clades. Co-infection may be more common than detected given our sampling strategy (*i.e.*, propagation and sequencing of only a few isolates per individual and only a portion of a single gene).

**Figure 3.**
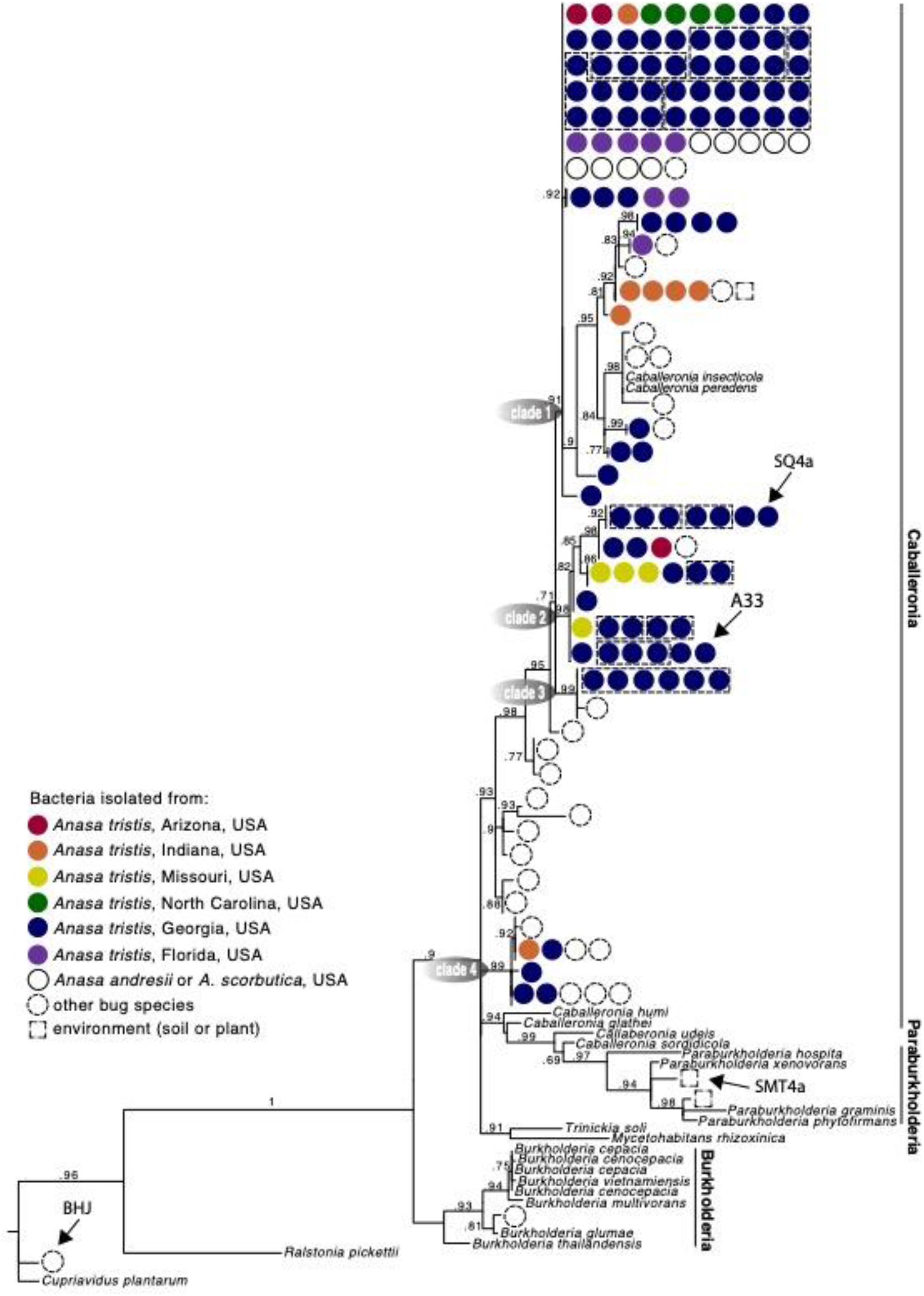
**Phylogeny of *Caballeronia* isolated from squash bug crypts, including symbionts of other true bugs species and other Burkholderiaceae**, based on 854 base pair alignment of 16s rRNA. Dashed lines group isolates within a clade that were isolated from the same individual. Detailed taxa information is provided in Supplemental Figure 1. Arrows point to isolates used for fitness assays, and clades are mentioned in main text.

### 3.3 Symbiont Transmission

In order to determine how *Caballeronia* are transmitted and acquired, we inoculated a generation of *A. tristis* with GFP-labeled *Caballeronia* sp. SQ4a and let them mate freely in a non-sterile environment with plants and soil. We then screened their offspring for the presence of the fluorescent *Caballeronia*. No fluorescent *Caballeronia* were found on or within eggs or in first instar nymphs. Fluorescent *Caballeronia* were recovered from 50 – 75% of second, third, fourth, and fifth instar nymphs, as well as adult offspring (Fig. 4). Using a fluorescent microscope, visualization of whole crypts dissected from adult offspring indicated that 11 of 15 crypts examined were colonized by GFP-labeled bacteria; five of six examined crypts from fifth instars were colonized. Given the substantial fitness benefits of *Caballeronia* association (see below), we presume that many individuals that made it to later instars without GFP-labeled *Caballeronia* may have picked up different strains from the plant or soil, neither of which were sterile, as crypts of these individuals often contained other, non-GFP-labeled bacteria that looked morphologically similar to *Caballeronia*.

**Figure 4.**
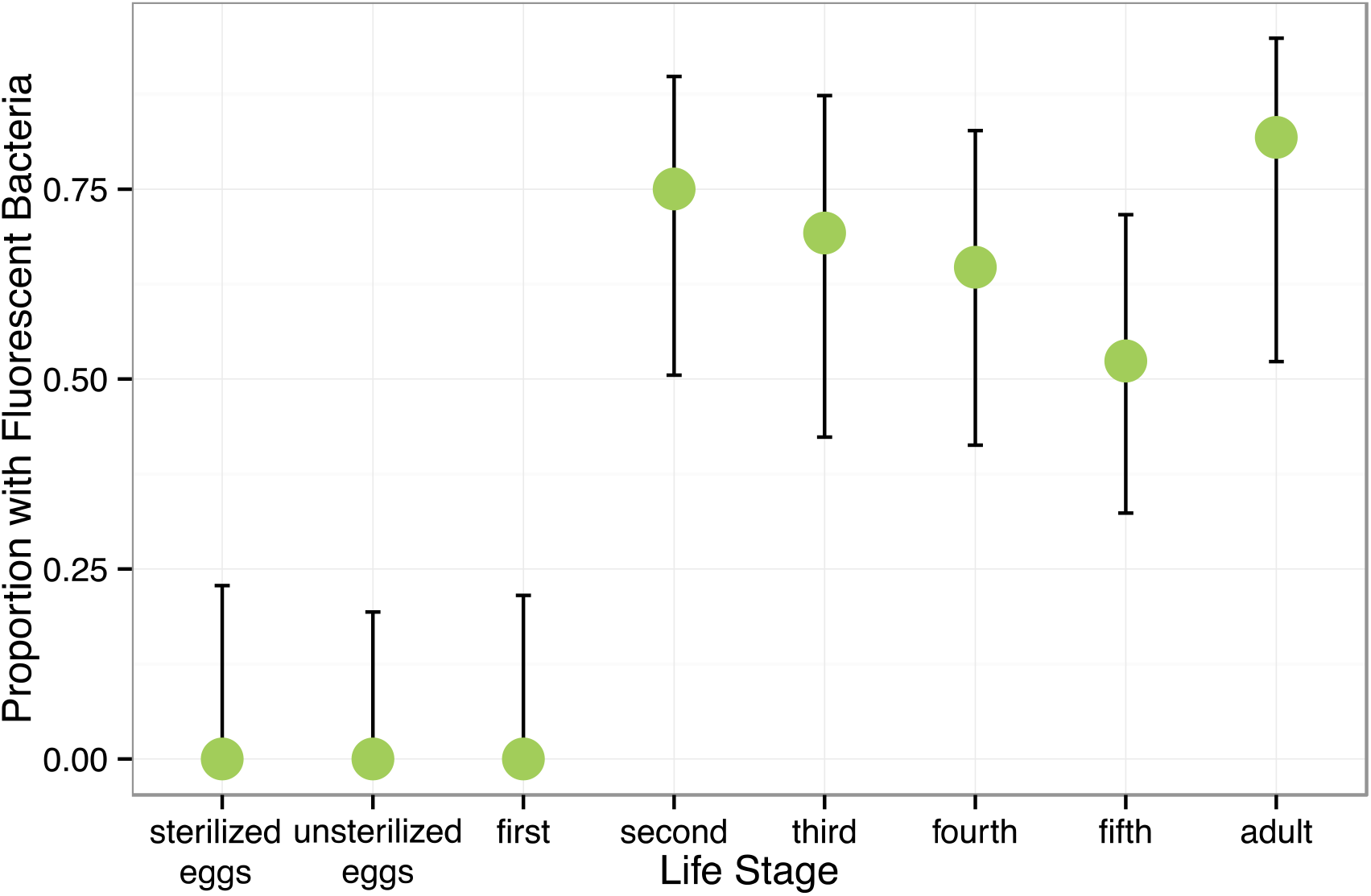
Proportion of offspring of different life stages harboring *Caballeronia* sp. SQ4a when reared on plants with parents previously inoculated with *Caballeronia* sp. SQ4a. Error bars are binomial confidence intervals. Sample sizes are 13 sets of two sterilized eggs, 16 sets of two unsterilized eggs, 14 first instar nymphs, 18 second instar nymphs, 14 third instar nymphs, 17 fourth instar nymphs, 23 fifth instar nymphs and 11 adult offspring.

### 3.4 Symbiont Uptake and Colonization

Despite exposure to large numbers of bacteria using three different feeding methods, only 5 of 27 first instars exposed to *Caballeronia* established *Caballeronia* infections. In comparison 20 of 20 second instar nymphs fed using similar methods established *Caballeronia* infections (Fig. 5). These results are consistent with the transmission experiment, above, in which we found that colonization increased dramatically after bugs molted to the second instar.

**Figure 5.**
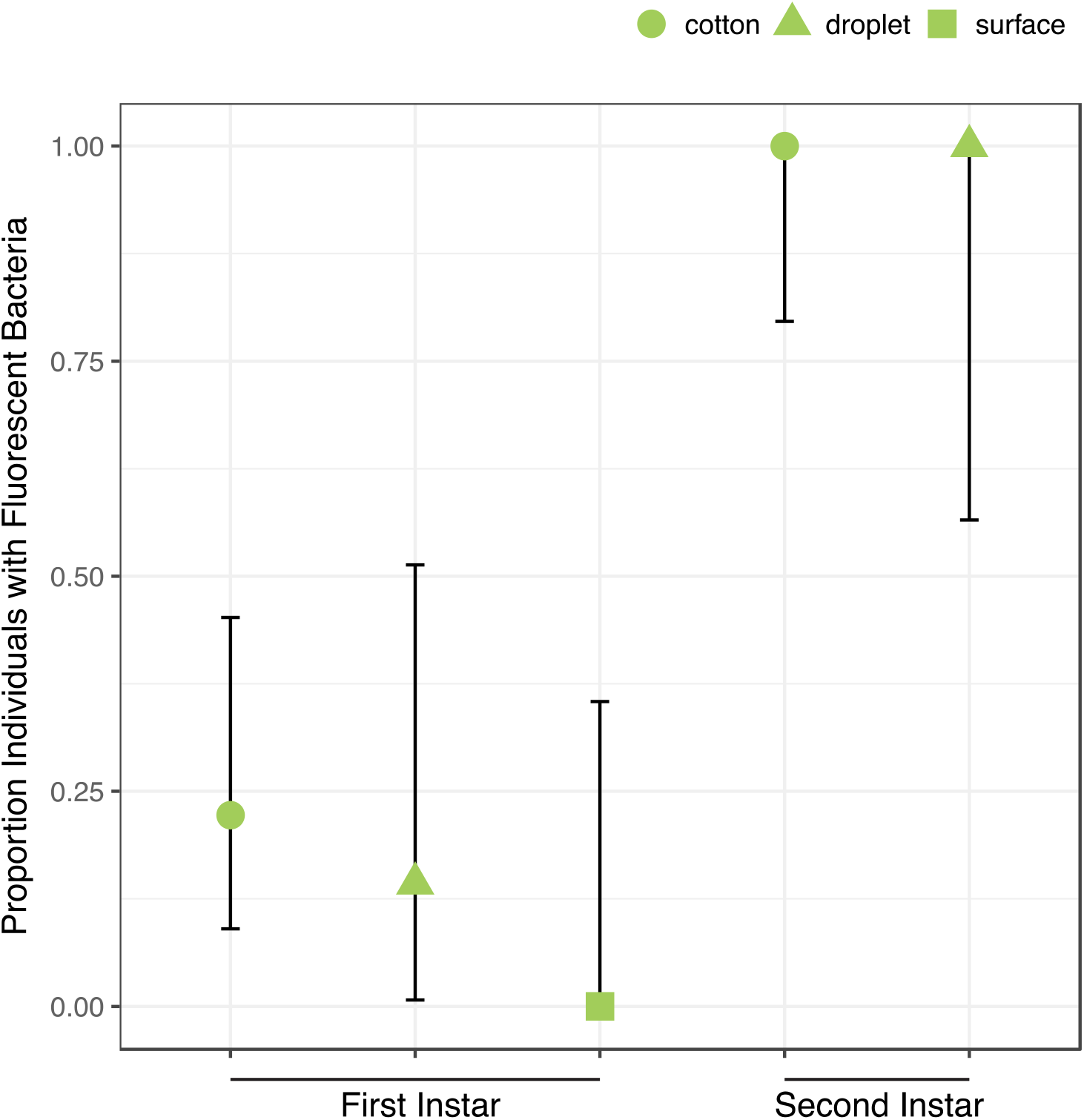
Acquisition of *Caballeronia* is largely constrained to later instars. Using three feeding methods (“cotton”, “droplet”, “surface”) in which squash bugs were exposed to solutions containing high concentrations of *Caballeronia* sp. SQ4a, first instars (N = 27) only rarely established infections. In comparison, when second instars (N = 20) were fed using two of the three methods, they always established the symbiosis. Error bars are binomial confidence intervals.

Upon reaching adulthood, both males and females were colonized with a large population of *Caballeronia* sp. SQ4a in their midgut crypts, with populations averaging 3.46 × 10^5^ CFUs per adult (Fig. S3). Though we have observed that their crypts do often appear to be bigger, females did not harbor significantly more *Caballeronia* sp. SQ4a than males (T-test on log-transformed data: *t*_*df*=18.94_ = 1.69, *P* = 0.11).

### 3.5 Fitness Benefits of Symbiosis with a Focal Symbiont for Host when Reared on Plants

In our first assay of the fitness effects of symbiosis with *Caballeronia* sp. SQ4a, during which individuals were reared on plants, survival to adulthood was significantly higher for insects inoculated with *Caballeronia* sp. SQ4a than for control insects inoculated with sterile water (Χ^2^_df=1_= 11.71, *P* < 0.001; HR = 0.35, 95% CI = 1.49 to 5.48; Fig. 6a). The proportion of insects surviving and molting to the third instar in *Caballeronia* sp. SQ4a replicates was not significantly different than those in water (control) replicates (F_1,10_ = 2.79, *P* = 0.13, odds ratio = 0.30, 95% C.I.= −2.87 to 0.20). Insects inoculated with *Caballeronia* sp. SQ4a, however, were significantly more likely to survive and molt to the fourth instar (F_1,10_ = 9.32, *P* = 0.01, odds ratio = 0.11, 95% C.I.= −3.78 to −0.75), fifth instar (F_1,10_ = 26.65, *P* < 0.001, odds ratio = 0.05, 95% C.I. = −4.25 to −1.72) and into adults (F_1,10_ = 121.88, *P* < 0.001, odds ratio = 0.001, 95% C.I. = −8.26 to −3.72; Fig 6b). Indeed, no control insects survived and molted into adults. For those insects surviving to the third instar, development time from hatch to third instar was significantly shorter for those inoculated with *Caballeronia* sp. SQ4a than those inoculated with water (W = 247.00, *P* < 0.001, 95% CI = −3.00 to −1.00; Fig 6c). This was also true for those surviving to the fourth (W = 52.50, *P* < 0.001, 95% CI = −11.00 to −8.99) and fifth instars (W = 28.00, *P* < 0.001, 95% CI −15.99 to −11.00; Fig 6c). Differences in development time from hatch to adult could not be assessed, as no water-inoculated controls survived and molted into adults in this experiment.

**Figure 6.**
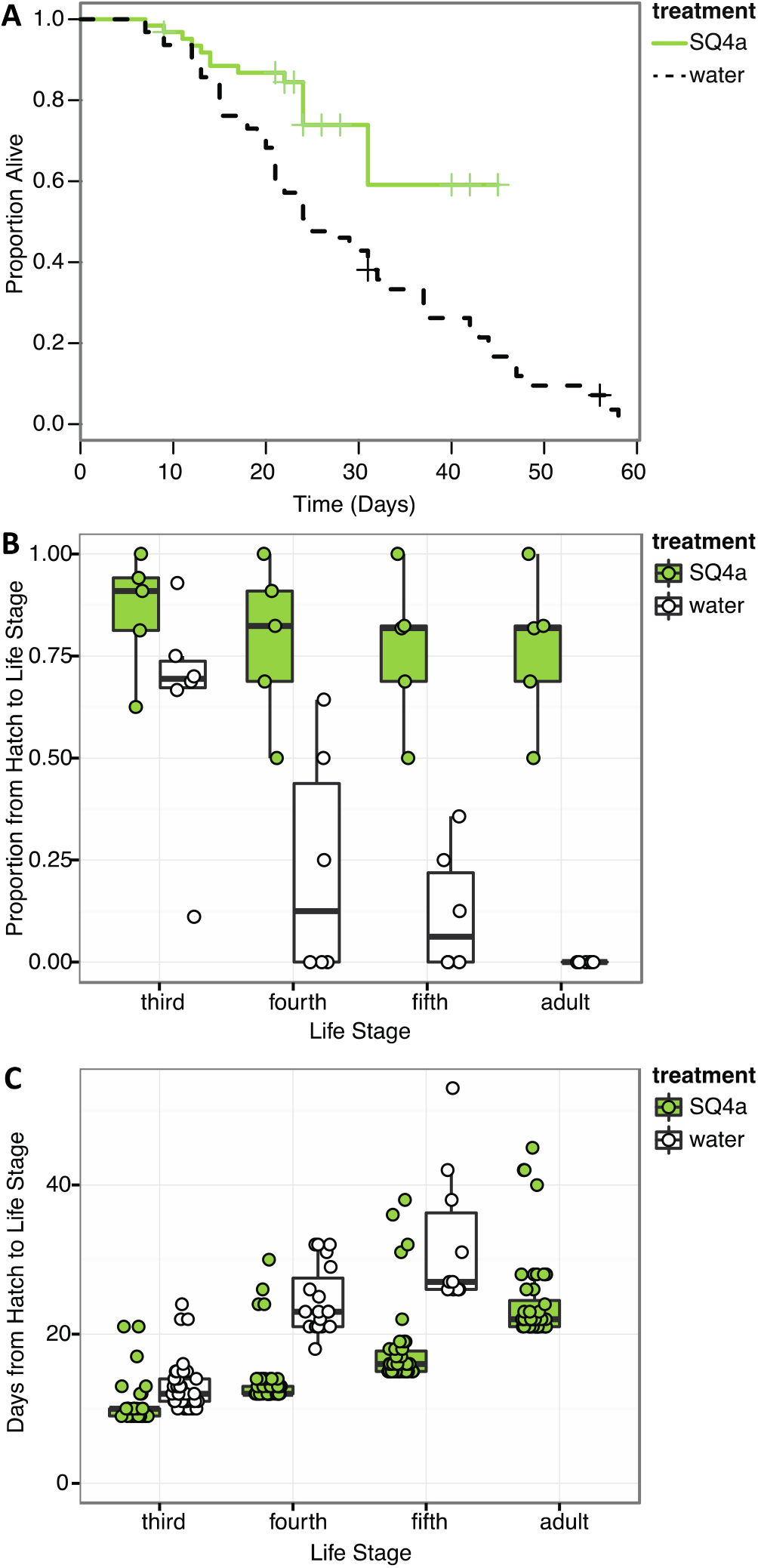
*Caballeronia* sp. SQ4a association increases survival and decreases development time when squash bugs are reared on plants. **(A)** Kaplan-Meier survival curves for the *Caballeronia* inoculation treatment using *Caballeronia* sp. SQ4a (N = 62) and the water (control) (N = 63) treatment. Data is censored at the time individuals reached adulthood or, in a few cases, when individuals escaped from cages (indicated by crosses) **(B)** Proportion of individuals in each tent surviving from hatch to each subsequent life stage. Proportions for each tent are overlaid on box-whisker plots. **(C)** Development time from hatch to each instar. Points, overlaid on box-whisker plots, indicate estimated development time for each individual that made it to that life stage. In this experiment, no water-inoculated (control) individuals survived to adulthood.

### 3.6 Fitness Benefits of Symbiosis with a Focal Symbiont for Host when Reared on Plants versus Fruits

In our second assay of the fitness effects of symbiosis with *Caballeronia* sp. SQ4a, during which individuals were reared on either plants or fruits, survival to adulthood was higher for *Caballeronia*-infected insects than non-infected ones, regardless of the feeding substrate (symbiosis: X^2^_df=1_ = 49.67, *P* < 0.001, HR = 0.18, 95% C.I.=2.77 to 11.1; substrate: X^2^_df=1_ = 1.74, *P* = 0.19, HR = 1.35, 95% C.I. = 0.31 to 1.79; interaction: X^2^_df=1_ = 0.01, *P* = 0.9, HR= 1.07, 95% C.I. = 0.34 to 2.6; Fig. 7a). We did not find a clear effect of substrate on the proportion of insects that survived from hatching to each instar (Fig. 7b). While a higher proportion of *Caballeronia*-infected nymphs fed on plants versus fruits made it from hatching to third instar (F_1,15_ = 9.71, *P* < 0.01, odds ratio = 2.1, 95% C.I.= −0.1 to 1.62) and to adulthood (F_1,15_ = 57.62, *P* = 0.04, odds ratio = 3.31, 95% C.I. = −0.14 to 2.63), we did not observe such an effect on those that made it from hatching to fourth (F_1,15_ = 2.55, *P* = 0.14, odds ratio = 1.51, 95% C.I. = −0.38 to 1.22) or fifth instar (F_1,15_ = 3.1, *P* = 0.10, odds ratio = 1.47, 95% C.I. = −0.51 to 1.3). This pattern suggests that there might be factors driving developmental differences between nymphs feeding on fruit or plant that were unaccounted for (*i.e.*, humidity, cage size, crowding, etc.). However, *Caballeronia* infection significantly increased the proportion of nymphs that made it from hatching to each developmental stage (third: F_1,15_ = 74.67, *P* < 0.001, odds ratio = 0.43, 95% C.I. = −1.53 to −0.16; fourth: F_1,15_ = 147.21, *P* < 0.001, odds ratio = 0.03, 95% C.I. = −4.72 to −2.69; fifth: F_1,15_ = 107.65, *P* < 0.001, odds ratio = 0.01, 95% C.I. = −7.00 to −3.14; and adulthood: F_1,15_ = 28.14, *P* < 0.01, odds ratio = 0.05, 95% C.I.= −8.39 to −0.71). The proportion of insects that made it to third instar was also affected by the interaction between substrate and *Caballeronia* infection (F_1,15_ = 20.69, *P* < 0.001, odds ratio = 0.09, 95% C.I. = −3.47 to −1.35).

**Figure 7.**
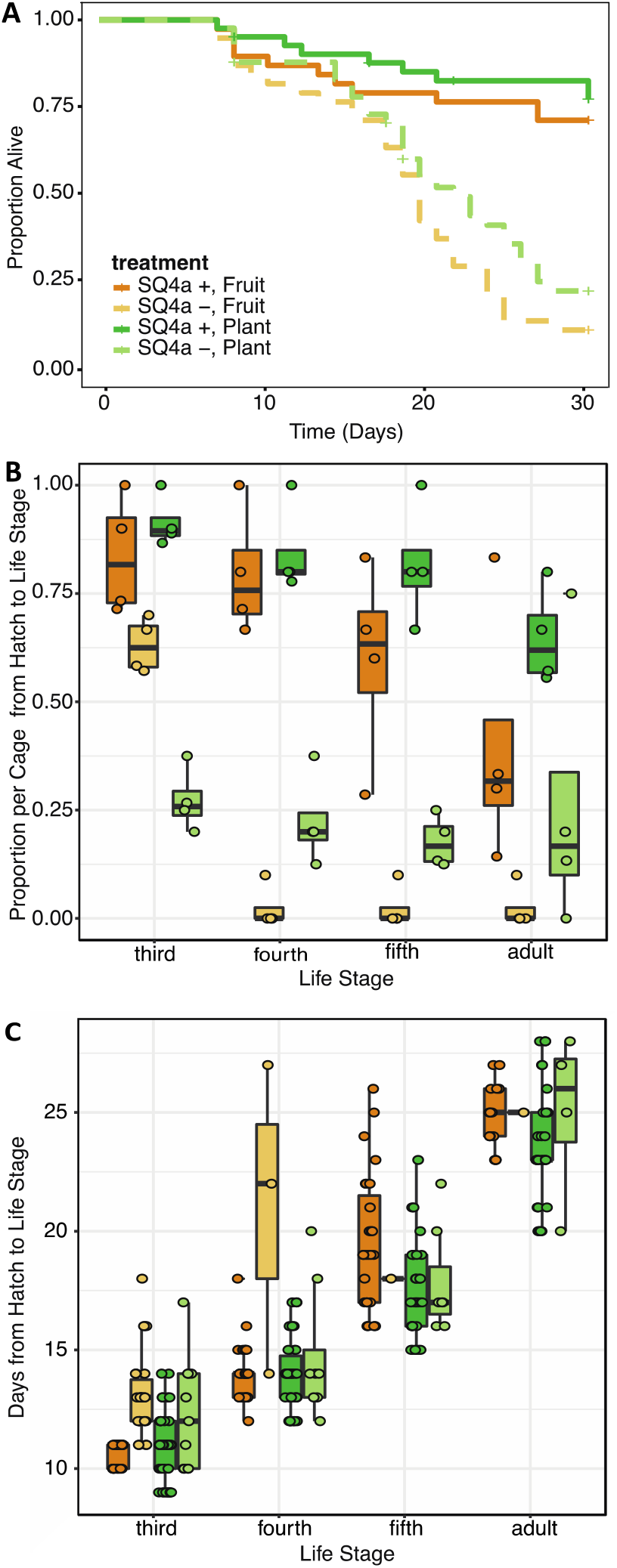
*Caballeronia* sp. SQ4a association provides similar survival and developmental benefits when squash bugs are reared on plants or fruits. **(A)** Kaplan-Meier survival curves for *Caballeronia*-inoculated (solid) and -uninoculated (dashed) treatments when bugs were reared on squash plants (green) and fruits (orange) (N = 41 individuals for each plant treatment and N = 38 for each fruit treatment). Data is censored at the time individuals reached adulthood or, in a few cases, when individuals escaped from cages (indicated by crosses) **(B)** Proportion of individuals in each cage surviving from hatch to each subsequent life stage (color scheme as in (A)). Proportions for each cage are overlaid on box-whisker plots. **(C)** Development time from hatch to each instar (color scheme as in (A)). Points, overlaid on box-whisker plots, indicate estimated development time for each individual that made it to that life stage.

Feeding substrate had no effect on the time nymphs took to develop to any life stage except fifth instar, in which case nymphs developed faster on plants versus on fruits (third: F_1,106_ 0.96, *P* = 0.33, OR = 1.03, 95% C.I. = −0.04 to 0.10; fourth: F_1,73_ = 2.18, *P* = 0.14, OR = 0.99, 95% C.I. = −0.07 to 0.05; fifth: F_1,68_ = 8.84, p < 0.01, OR = 0.89, 95% C.I.= −0.18 to −0.04; and adult: F_1,49_ =10.07, *P* = 0.11, OR = 0.95, 95% C.I. = −0.11 to 0.01; Fig. 7c). *Caballeronia-*infected nymphs had shorter developmental time from hatching to third (F_1,106_ = 45.61, *P* < 0.001, OR = 1.23, 95% C.I. = 0.13 to 0.28), and fourth instar (F1,73 = 19.56, *P* < 0.001, OR = 1.50, 95% C.I.=0.27 to 0.53). After fourth instar, infection status no longer impacted developmental time (fifth instar: F_1,68_ = 0.66, p = 0.42, OR = 0.92, 95% C.I. = −0.36 to 0.18; and adulthood: F_1,49_ = 9.75, *P* = 0.61, OR = 1.06, 95% C.I. = −0.15 to 0.27); this is likely driven by the fact that very few individuals without symbionts molted to these life stages.

### 3.7 Variation in Fitness Benefits Conferred by Alternative Bacteria

Compared to the two fitness experiments described above, survival to adulthood was much higher in this third fitness experiment. Here, 91% of *Caballeronia*-inoculated individuals survived to adulthood, and 84% of control insects survived to adulthood. In contrast, in our first fitness experiment, 77% of *Caballeronia*-inoculated individuals survived to adulthood, and no control insects inoculated with sterile water survived to adulthood. We hypothesize that these differences are due to optimal rearing conditions for this experiment, in which bugs were at lower population density than the other experiment on fruit, bugs were transferred to new fruit each day, and rearing chambers were cleaned rigorously each day. Under these conditions, there was no significant difference in survival across the five treatments (χ^2^_df=4_ = 8.01, *P* = 0.09; Fig. 8a). In terms of proportion of individuals surviving to each instar, all 353 experimental individuals survived to third instar, and most (349 of 353) individuals survived from hatch to fourth instar. There was a significant effect of treatment on the proportion of individuals that made it from hatch to later life stages (fifth: F_4,87_ = 4.63, *P* < 0.01; adult: χ^2^_df=4_ = 85.99, *P* < 0.001; Fig. 8b), though survival to these later life stages too was high.

**Figure 8.**
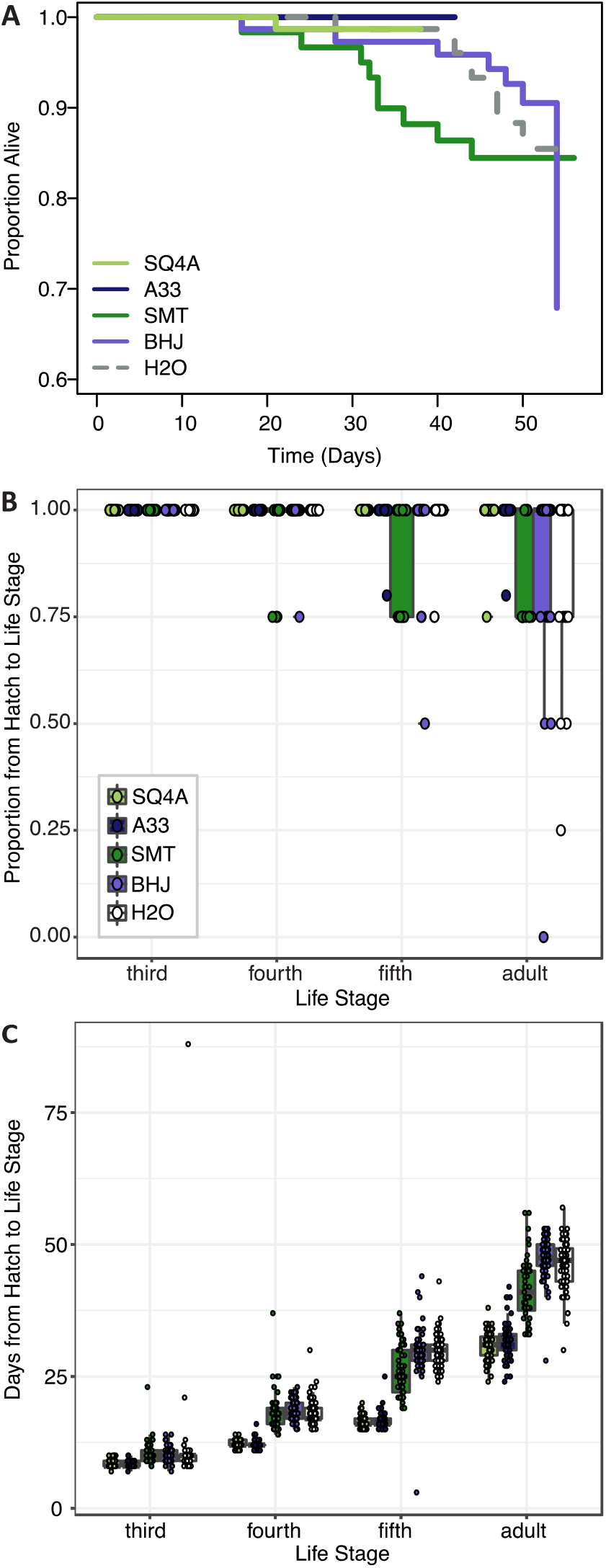
Bacterial strains provide different levels of fitness benefit. Squash bugs were fed either one of two strains of *Caballeronia* (SQ4a, A33), a strain of *Paraburkholderia* (SMT), a strain of *Cupriavidus* (BHJ), or water (H20) as an un-inoculated control. **(A)** Kaplan-Meier survival curves for bacteria-inoculated (solid lines) and un-inoculated (dashed) treatments. Data is censored at the time individuals reached adulthood. **(B)** Proportion of individuals in each cage surviving from hatch to each subsequent life stage. Proportions for each cage are overlaid on box-whisker plots. **(C)** Development time from hatch to each instar (legend in (B)). Points, overlaid on box-whisker plots, indicate estimated development time for each individual that made it to that life stage.

There were significant differences in development time based on treatment (Fig. 8c; hatch to third instar: F_4,351_ = 5.30, *P* < 0.01; hatch to fourth instar: F_4,346_ = 202.70, *P* < 0.001; hatch to fifth instar: F_4,336_ = 356.33, *P* < 0.001; hatch to adult: F_4,313_ = 236.88, *P* < 0.001). Notably, development time for individuals inoculated with either *Caballeronia* sp. SQ4a or A33 was consistently significantly shorter than development time for individuals not inoculated with *Caballeronia*, while development time for individuals fed BHJ was not different than that of those not given a symbiont. SMT-inoculated individuals developed significantly faster to the adult stage than individuals not inoculated with *Caballeronia*, but they developed significantly more slowly than individuals inoculated with SQ4a and A33 (Supplemental Table 4).

Both bacterial treatment and sex significantly influenced adult body size, as measured by the width of the pronotum (Fig. 9a; treatment: F_4,182_ = 16.05, *P* < 0.001; sex: F_1,182_ = 311.38, *P* < 0.001). There was no significant interaction of bacterial treatment and sex (F_4,182_ = 0.84, *P* = 0.5). Focusing on bacterial treatments, individuals inoculated with SQ4a and A33 were significantly bigger than individuals fed BHJ, SMT or water (Table S5). Qualitatively, sex and treatment had similar impacts on other measurements of body size (*i.e.*, body length, scutellum width, tibia length), as many measurements of *Anasa* spp. body size are positively correlated (data not shown). In terms of weight, just after molting to the adult stage, females were significantly heavier than males (Fig. 9b; F_1,251_ = 192.58, *P* < 0.001). Bacterial treatment did not have a significant impact on adult weight (F_4,251_ = 1.09, *P* = 0.36), nor was there a significant interaction between sex and treatment (F_4,251_ = 0.43, *P* = 0.79).

**Figure 9.**
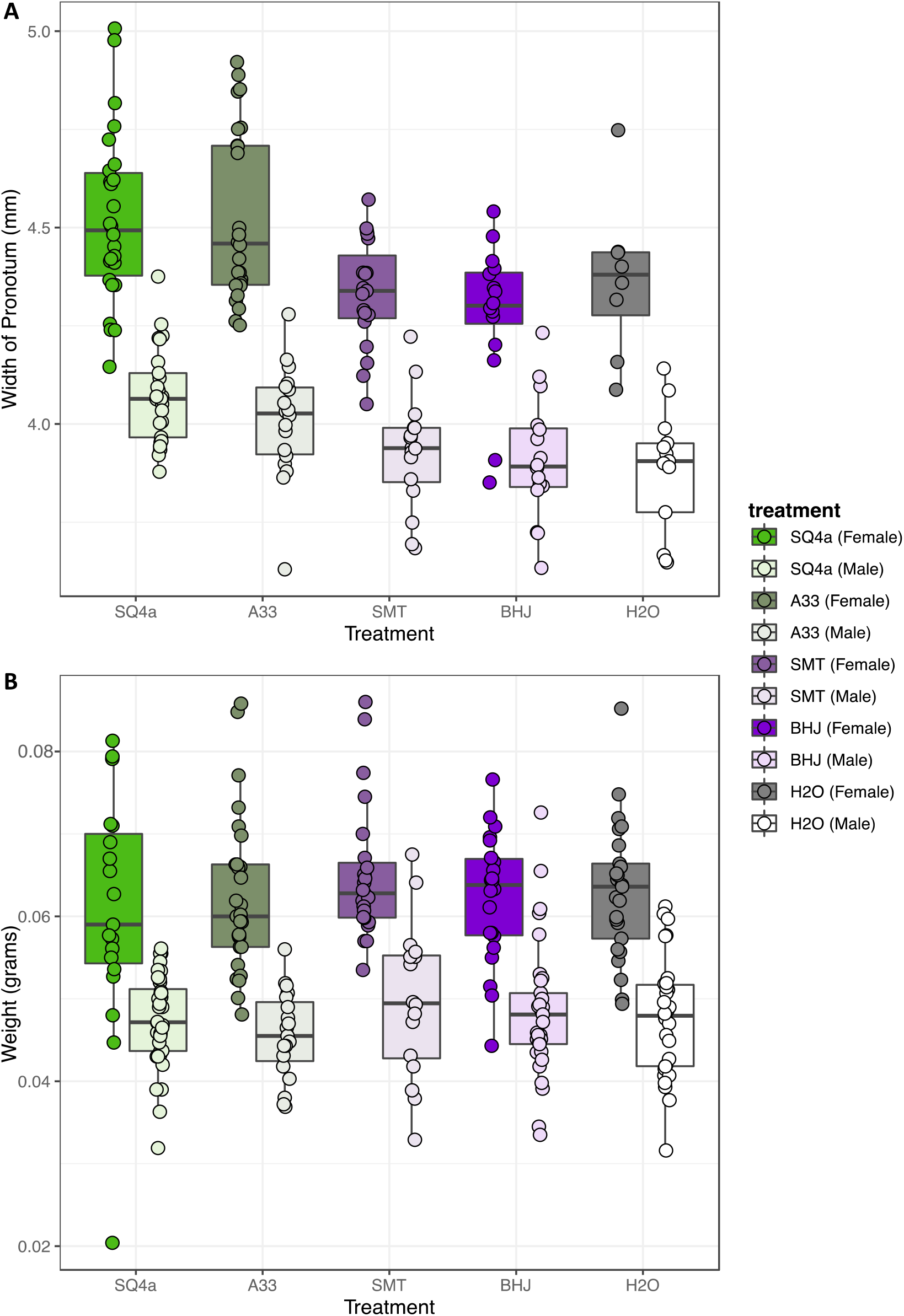
Bacterial treatment significantly influences size but not weight of adult squash bugs. **(A)** Based on the width of the pronotum, females were larger than males, and, overall bugs inoculated with SQ4a and A33 were significantly bigger than individuals fed SMT, BHJ and water. **(B)** Females were heavier than males, but there was no overall effect of bacterial treatment on adult wet weight.

### 3.8 Colonization by Alternative Bacteria

The four strains varied in terms of their final loads of GFP-labeled bacteria within the bugs’ crypts (Fig. 10). Estimated CFUs of SMT in crypts (mean CFUs) were higher than estimated CFUs of A33 and SQ4a (means of 1.31 × 10^6^ and 3.46 × 10^5^ CFUs, respectively); all adults screened for these strains were colonized by GFP-labeled bacteria only. BHJ, which is in the genus *Cupriavidus*, exhibited extremely low colonization (mean 1.23 × 10^2^ CFUs) and was recovered from only 3 of 15 sampled adults. For BHJ, colonization should be interpreted with caution; it is possible that recovered GFP-labeled bacteria were not in the crypts but were contaminants from nearby sections of the gut. No GFP-labeled bacteria were isolated from water-fed controls, and non-GFP labeled bacteria, which were not quantified, never reached high titers. Considering only the three strains that consistently colonized the crypts, differences in final symbiont load were significantly impacted by the bacterial strain (F_2,51_ = 26.17, *P* < 0.001), and to a lesser extent by sex (F_1,51_ = 6.16, *P* = 0.02); there was no significant interaction between bacterial strain and sex (F_2,51_ = 1.56, *P* = 0.22).

**Figure 10.**
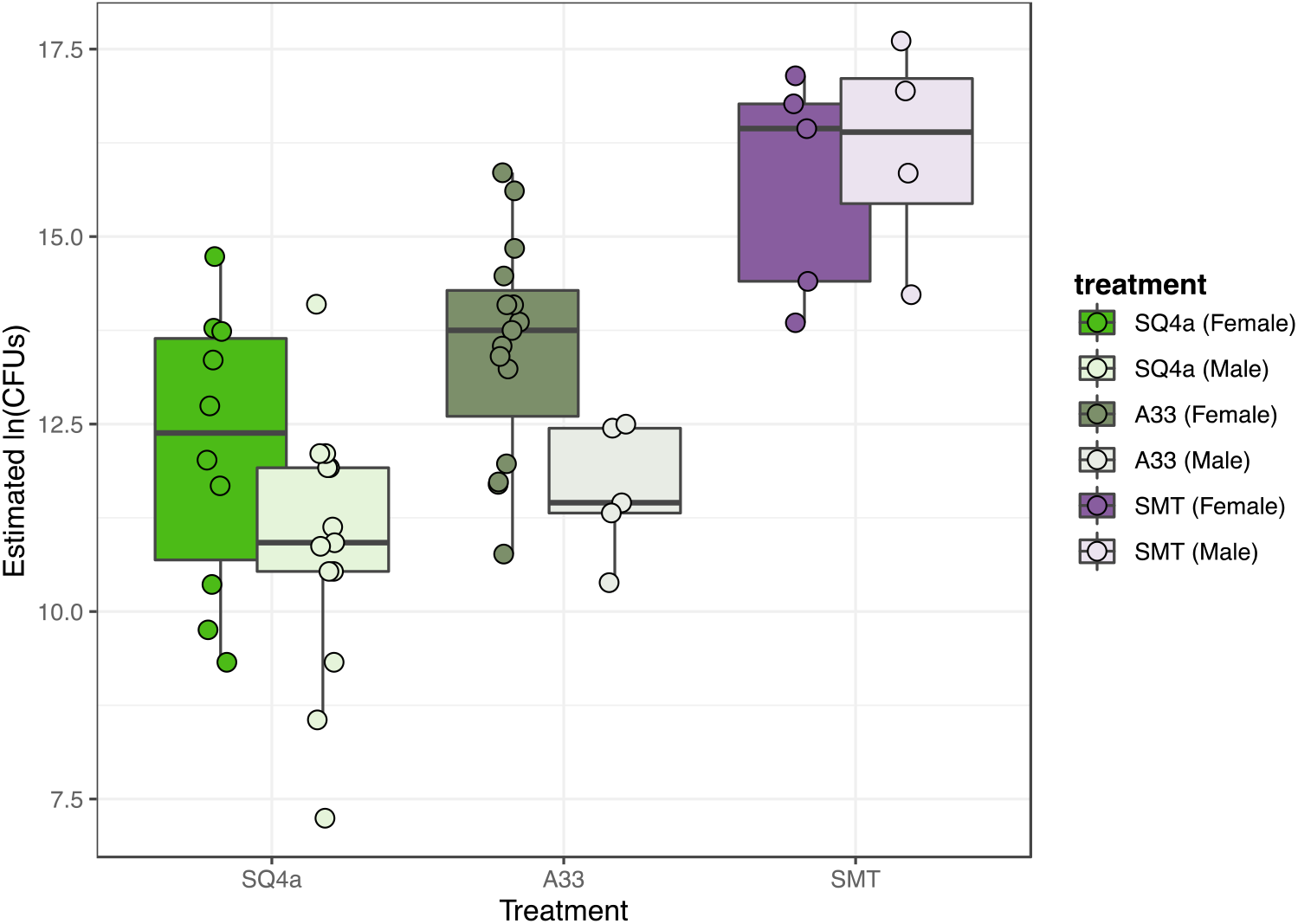
Estimated population sizes of alternative Burkholderiaceae strains in adult crypts. The populations sizes in crypts of the three strains that established in adult bugs differed significantly, with SMT, the *Paraburkholderia* sp. strain that provided less fitness benefits (Figure 9), establishing the largest populations.

## 4 Discussion

The prevalence of *Caballeronia* infection and the dominance of *Caballeronia* in *A. tristis’* midgut crypt microbial communities suggests that *Caballeronia* is a symbiont of *A. tristis*, as it is in many bug species (Boucias et al., 2012; Garcia et al., 2014; Gordon et al., 2016; Itoh et al., 2014; Kikuchi et al., 2005, 2011a; Ohbayashi et al., 2019b; Olivier-Espejel et al., 2011; Ravenscraft et al., 2020; Sudakaran et al., 2015; Takeshita et al., 2015; Xu et al., 2016a). Experiments varying both the environment and the symbiont strain indicate that the bacteria, which are environmentally acquired, can both increase survival to adulthood and decrease development time, confirming the benefit of *Caballeronia*-association for these pests. Both sequencing of *Anasa*-associated *Caballeronia* and fitness experiments with alternative strains highlight genetic and phenotypic diversity within the system that could have important ecological and evolutionary consequences for the hosts and their symbionts.

Though the bacterial community within a crypt appears to often be dominated by a single *Caballeronia* strain, there does appear to be occasional co-infections of crypts. Both the culture-independent and culture-dependent approaches indicate that bacteria of other genera can infect the crypts, though at low levels relative to *Caballeronia*. Furthermore, based on culture-dependent sequencing, which may be limited in its ability to capture co-infections if some strains are recalcitrant to *in vitro* culturing, a few individuals were determined to be co-infected with multiple *Caballeronia* strains. The low frequency of co-infections is similar to some *Caballeronia-*associated species (e.g. *Riptortus pedestris*, *Blissus insularis*) that are rarely co-infected with multiple strains (Boucias et al., 2012). Broad-headed bugs (*Alydus* spp.) and stilt bugs (*Jalysus* spp.), however, can be co-infected with multiple *Caballeronia* strains at higher frequencies (Garcia et al., 2014; Ravenscraft et al., 2020). Though some of this variation is likely due to sampling differences, it could also reflect differences in host ecology or physiology that alter the likelihood of co-infection or differences in the ability of the bacteria to outcompete one another (Itoh et al., 2019). Further sampling and experiments will be necessary to understand the environmental and evolutionary factors that contribute to differences in co-infection prevalence across species. Future experiments are also needed to determine the impact of crypt co-infection, whether by bacteria of multiple genera or by multiple strains of *Caballeronia*, on host and symbiont fitness.

An effective and efficient acquisition mechanism is necessary for hosts to maintain a microbial symbiont. Our rearing experiments indicate that *Caballeronia* sp. SQ4a is not directly passed on to offspring internally or on egg surfaces, which are common routes of vertical transmission, particularly for obligate symbionts, in insects (Salem et al., 2015). Instead, squash bugs acquire *Caballeronia* expelled from adults via the environment during, and possibly after, the second instar. Environmental acquisition is also largely confined to the second instar in bean bugs (Kikuchi et al., 2011b), indicating that there may be a developmentally confined signal or behavior required for symbiont acquisition in the Coreoidea superfamily of insects, or that there may be physiological differences between instars that constrain symbiont acquisition to a narrow window. In experiments in which we know that first instars probed liquid containing live bacterial cells (*i.e.*, droplet feeding method), we rarely observed establishment, suggesting possible physiological constraints on symbiont establishment.

In our transmission experiment, which involved tracking of the acquisition of a fluorescently-labeled *Caballeronia* strain, it is unclear from where in the environment the nymphs acquired the bacteria. At the onset, adult squash bugs were the only source of labeled *Caballeronia* in our rearing experiments, but it is unlikely that nymphs are picking up these bacteria through direct contact with the adults, as such contact appears rare. We hypothesize that *Caballeronia* are transmitted from parents or other individuals through two potential environmental routes. *Caballeronia* could be excreted onto plant surfaces or soil with waste products, or they could be transmitted to the surface or internal tissue of the plant as the adults feed, which occurs for other insect symbionts (Gonella et al., 2015; Xu et al., 2016b). Further research will be required to tease apart these alternative routes of environmental acquisition and whether mechanisms of transmission differ based on symbiont strain. Regardless, the fact that the bacteria expelled from adults can be transmitted to the next generation of hosts is consistent with the fact that most individuals we sampled in the field are associated with closely-related strains, and suggests a possible combination of horizontal transmission and indirect vertical transmission from parents through an environmental route. Such a transmission route, in which symbionts are escaping into the environment where they are readily picked up by the next generation, may limit some of the negative consequences of vertical transmission, in which population structure imposed by strict host association, coupled with a bottleneck every generation, can lead to symbiont degeneration (O’Fallon, 2008). Environmental acquisition, however, does have risks, as bugs could not acquire a symbiont, or could acquire a less beneficial strain.

Indeed, we experimentally demonstrated the possibility for individuals to pick up strains that confer different levels of fitness benefits. In a comparison of insect fitness upon inoculation with one of four different environmental bacterial strains, we find substantial differences in a number of life history traits, including likelihood of molting to adulthood, development time and adult body size. The ability of alternative strains to colonize and provide differential fitness benefits to their hosts has also been shown for *R. pedestris* (Itoh et al., 2019), the best studied *Caballeronia*-associated true bug species. The variation in benefits may be driven by the metabolic benefits that the bacteria can provide their hosts. Genomic and transcriptomic investigations indicate that the symbiont of *R. pedestris*, *B. insecticola*, is involved in carbon, sulfur and nitrogen metabolism. The bacterium also produces all essential amino acids and B vitamins (Ohbayashi et al., 2019a). Whether these are the benefits for *A. tristis* remains to be determined, but the breadth of benefits provided for *R. pedestris*, along with the many bacterial phenotypes that appear necessary for successful host colonization (Kim et al., 2013a, 2013b, 2014; Kinosita et al., 2017), open up the possibility for substantial variation across symbiont strains, particularly if selection acts on these bacteria such that some functions are lost due to relaxed selection or purifying selection when the bacteria are outside of the host. Future research should investigate the bacterial and host factors that underlie variation in symbiotic benefit.

Considering the symbiosis from the perspective of the symbiont, bacterial fitness, as measured by colonization of bug crypts, also varied across bacterial strains used in our study. While the two *Caballeronia* strains (SQ4a, A33) colonized bugs to a similar level, SMT4a, a *Paraburkholderia* strain that was less beneficial for the bugs, established significantly larger population sizes in the bugs. Whether these larger populations are costly for the bugs is unclear. It is also unknown whether reaching a larger population size in hosts would increase the likelihood of the bacteria being transmitted to the environment or to other individuals, which may be a better proxy for considering the impacts of host colonization on bacterial fitness. The variation seen across the few strains screened here suggests that the *A. tristis* system could be used to explore this question.

### 4.1 Conclusions

In this study, we demonstrate that bacterial communities within midgut crypts of *Anasa tristis* are dominated by *Caballeronia* symbionts. Fitness assays on two different substrates suggest that *Caballeronia* is an essential symbiont of these squash bugs in natural populations. Given that the presence of the symbiont is only detected after molting to the second instar, *A. tristis* likely acquires the symbiont from the environment. Environmental acquisition, coupled with the fact that bacteria strains vary in the level of fitness benefits that they provide to their hosts, provides an opportunity for these insect pests to acquire bacteria that have alternative phenotypic effects. This opens up the possibility to manipulate the symbiosis in agricultural settings to alter the fitness and/or vector competence of this pest insect and vector of plant disease.

## 5 Data Availability Statement

All experimental datasets and code for statistical analyses will be made available in Dryad upon acceptance. The nucleotide sequence data are available in the DDBJ/EMBL/GenBank database under accession numbers: KT259132 - KT259191, KX239751 - KX239768, MH636869 - MH636872 and MZ264232-MZ264276.

## 6 Author Contributions

TSA, GPF, JRG, AB and NMG designed experiments. TSA, GPF, JRG, TA, AB and KSS conducted experiments. TSA, GPF, JRG, AB and NMG analyzed data. All authors contributed to writing and approved the final version of the manuscript.

## 7 Funding

Research was supported by National Science Foundation grant IOS-1149829 to NMG, USDA NIFA 2019-67013-29371 to NMG, NSF Graduate Research Fellowships to JRG and KSS [grant number DGE-1444932] and a SSE RC Lewontin Early Graduate Research Award to KSS.

## 8 Acknowledgments

We thank the staff at Crystal Organic Farms, Oxford College Organic Farm, Oakhurst Garden, North DeKalb Community Garden, Front Field Farm, Ten Mother’s Farm, Farlow Farm, Merry Lea Sustainable Farm, DeCamp Gardens, and Woodland Gardens, as well as gardeners Sondra Stoy and Dorothy Barse. Special thanks to Nicolas Donck and Daniel Parsons for allowing us to collect samples and for always being supportive of our research. We thank Dr. Kent Shelby (University of Missouri), Cindy Goodman and Joseph Ringbauer for providing samples from the Biological Control of Insects Research Laboratory. We also thank Molly Hunter (Univeristy of Arizona), Scott Villa and Jason Chen for providing samples, and Sandra Mendiola for help with collecting. We thank Frank Stewart for providing access to a MiSeq, and Neha Sarode and Josh Parris for technical help with MiSeq assay design and sequencing.

## 9 Supplementary Materials

are available through the Bioarxiv site.

## 10 Conflict of Interest

The authors declare that the research was conducted in the absence of any commercial or financial relationships that could be construed as a potential conflict of interest.

## Notes

### Competing Interest Statement

The authors have declared no competing interest.

