## Supplementary Materials for "The importance of environmentally-acquired bacterial symbionts for the squash bug (*Anasa tristis*), a significant agricultural pest"

**Supplemental Table 1:**

| Sample | No. Raw Reads | No. Reads in Final Analysis | Read Length in Final Analysis | | |
|---|---|---|---|---|---|
| | | | Min. | Avg. | Max. |
| SB33-m1 | 127,976 | 101,129 | | | |
| SB34-m1 | 135,311 | 99,738 | | | |
| SB35-m1 | 129,943 | 105,604 | | | |
| SB36-m1 | 148,449 | 110,127 | | | |
| SB32-m4 | 131,497 | 106,878 | | | |
| SB33-m4 | 157,265 | 121,165 | | | |
| SB34-m4 | 95,709 | 75,902 | | | |
| SB35-m4 | 78,500 | 61,120 | | | |
| SB36-m4 | 123,032 | 92,575 | | | |
| SB74-wg | 89,147 | 70,057 | | | |
| SB80-wg | 203,580 | 156,361 | | | |
| SB84-wg | 138,314 | 105,484 | | | |
| SB85-wg | 127,780 | 101,425 | | | |
| SB86-wg | 142,729 | 109,106 | | | |
| **Total:** | **1,829,232** | **1,416,671** | | | |

**Supplemental Table 1. *Caballeronia* spp. bacteria isolated from *Anasa tristis.*** Location abbreviations indicate: BCI - BCI Research Laboratory, Columbia, Missouri; CRY - Crystal Organic Farms, Newborn, Georgia; DEC - DeCamp Gardens, Albion, Indiana; FFA - Farlow Farm, Archdale, Indiana; FFF - Front Field Farm, Winterville, Georgia; MGA - Mission Garden, Tucson, Arizona; MLF - Merry Lea Farm, Albion, Indiana; NDG - North Dekalb Garden, Atlanta, Georgia; OAK - Oakhurst Community Garden, Decatur, Georgia; OXF - Oxford Farm, Oxford, Georgia; SBG - Stoy/Barse Gardens, Auburn, Indiana; TMF - Ten Mothers Farm, Hillsborough, North Carolina; UFL – University of Florida Garden, Gainesville, Georgia; and WOG - Woodland Gardens, Atlanta, Georgia. All were collected within the United States. Classification and % match to genus are based on the Ribosomal Database Project (RDP) classifier.

| **Isolate Name** | **Individual** | **Life Stage** | **Tissue** | **State** | **Site** | **NCBI** | **Family** | ***Genus*** | **% Match to Genus** |
|---|---|---|---|---|---|---|---|---|---|
| 3_56_M4_b | 3.56 | 3 | M4 crypt | Georgia | CRY | KX239758 | Burkholderiaceae | *Caballeronia* | 100 |
| 3_56_M4_c | 3.56 | 3 | M4 crypt | Georgia | CRY | KX239759 | Burkholderiaceae | *Caballeronia* | 100 |
| 3_57_M4_a | 3.57 | 3 | M4 crypt | Georgia | CRY | KX239760 | Burkholderiaceae | *Caballeronia* | 100 |
| 3_57_M4_b | 3.57 | 3 | M4 crypt | Georgia | CRY | KX239761 | Burkholderiaceae | *Caballeronia* | 100 |
| 3_57_M4_c | 3.57 | 3 | M4 crypt | Georgia | CRY | KX239762 | Burkholderiaceae | *Caballeronia* | 100 |
| 4_50_M4_a | 4.5 | 4 | M4 crypt | Georgia | CRY | KX239751 | Burkholderiaceae | *Caballeronia* | 100 |
| 4_50_M4_c | 4.5 | 4 | M4 crypt | Georgia | CRY | KX239763 | Burkholderiaceae | *Caballeronia* | 100 |
| 4_51_M4_a | 4.51 | 4 | M4 crypt | Georgia | CRY | KX239764 | Burkholderiaceae | *Caballeronia* | 100 |
| 4_51_M4_b | 4.51 | 4 | M4 crypt | Georgia | CRY | KT259148 | Burkholderiaceae | *Caballeronia* | 100 |
| 4_51_M4_c | 4.51 | 4 | M4 crypt | Georgia | CRY | KT259151 | Burkholderiaceae | *Caballeronia* | 100 |
| 5_46_M4_a | 5.46 | 5 | M4 crypt | Georgia | CRY | KX239767 | Burkholderiaceae | *Caballeronia* | 100 |
| 5_46_M4_b | 5.46 | 5 | M4 crypt | Georgia | CRY | KX239768 | Burkholderiaceae | *Caballeronia* | 100 |
| A32_M4_a | A.32 | A | M4 crypt | Georgia | CRY | KX239752 | Burkholderiaceae | *Caballeronia* | 100 |
| A32_M4_b | A.32 | A | M4 crypt | Georgia | CRY | KX239753 | Burkholderiaceae | *Caballeronia* | 100 |
| A32_M4_c | A.32 | A | M4 crypt | Georgia | CRY | KX239754 | Burkholderiaceae | *Caballeronia* | 99 |
| A33_M4_a | A.33 | A | M4 crypt | Georgia | CRY | KX239755 | Burkholderiaceae | *Caballeronia* | 100 |
| A33_M4_c | A.33 | A | M4 crypt | Georgia | CRY | KX239756 | Burkholderiaceae | *Caballeronia* | 100 |
| ATUFL_F1_KS43 | ATUFL_F1 | A | M4 crypt | Florida | UFL | MZ264256 | Burkholderiaceae | *Caballeronia* | 100 |

| **Isolate Name** | **Individual** | **Life Stage** | **Tissue** | **State** | **Site** | **NCBI** | **Family** | ***Genus*** | **% Match to Genus** |
|---|---|---|---|---|---|---|---|---|---|
| ATUFL_F1_KS4A | ATUFL_F1 | A | M4 crypt | Florida | UFL | MZ264265 | Burkholderiaceae | *Caballeronia* | 100 |
| ATUFL_F2_KS42 | ATUFL_F2 | A | M4 crypt | Florida | UFL | MZ264255 | Burkholderiaceae | *Caballeronia* | 100 |
| ATUFL_F2_KS9A | ATUFL_F2 | A | M4 crypt | Florida | UFL | MZ264267 | Burkholderiaceae | *Caballeronia* | 100 |
| ATUFL_F3_KS10A | ATUFL_F3 | A | M4 crypt | Florida | UFL | MZ264276 | Burkholderiaceae | *Caballeronia* | 100 |
| ATUFL_M1_KS5A | ATUFL_M1 | A | M4 crypt | Florida | UFL | MZ264275 | Burkholderiaceae | *Caballeronia* | 100 |
| ATUFL_M2_KS44 | ATUFL_M2 | A | M4 crypt | Florida | UFL | MZ264257 | Burkholderiaceae | *Caballeronia* | 100 |
| AZ1_KS37 | AZ1 | A | M4 crypt | Arizona | MGA | MZ264251 | Burkholderiaceae | *Caballeronia* | 100 |
| AZ10_KS36 | AZ10 | A | M4 crypt | Arizona | MGA | MZ264250 | Burkholderiaceae | *Caballeronia* | 100 |
| AZ7_KS35 | AZ7 | A | M4 crypt | Arizona | MGA | MZ264272 | Burkholderiaceae | *Caballeronia* | 100 |
| GACF3 | GACF3 | A | M4 crypt | Gerogia | CRY | MZ264239 | Burkholderiaceae | *Caballeronia* | 100 |
| GACF4 | GACF4 | A | M4 crypt | Gerogia | CRY | MZ264273 | Burkholderiaceae | *Caballeronia* | 100 |
| GACF5 | GACF5 | A | M4 crypt | Gerogia | CRY | MZ264240 | Burkholderiaceae | *Caballeronia* | 100 |
| GAFFF1 | GAFFF1 | A | M4 crypt | Georgia | FFF | MZ264232 | Burkholderiaceae | *Caballeronia* | 100 |
| GAFFF2 | GAFFF2 | A | M4 crypt | Georgia | FFF | MZ264233 | Burkholderiaceae | *Caballeronia* | 100 |
| GAFFF3 | GAFFF3 | A | M4 crypt | Georgia | FFF | MZ264269 | Burkholderiaceae | *Caballeronia* | 100 |
| GAFFF5 | GAFFF5 | A | M4 crypt | Georgia | FFF | MZ264234 | Burkholderiaceae | *Caballeronia* | 100 |
| GAOx1 | GAOx1 | A | M4 crypt | Gerogia | OXF | MZ264271 | Burkholderiaceae | *Caballeronia* | 100 |
| GAWG1_1 | GAWG1_1 | A | M4 crypt | Gerogia | WOG | MZ264235 | Burkholderiaceae | *Caballeronia* | 100 |
| GAWG2_1 | GAWG2_1 | A | M4 crypt | Gerogia | WOG | MZ264237 | Burkholderiaceae | *Caballeronia* | 100 |
| GAWG2_2 | GAWG2_2 | A | M4 crypt | Gerogia | WOG | MZ264238 | Burkholderiaceae | *Caballeronia* | 100 |
| GAWG2_4 | GAWG2_4 | A | M4 crypt | Gerogia | WOG | MZ264274 | Burkholderiaceae | *Caballeronia* | 100 |
| INDec2 | INDec2 | A | M4 crypt | Indiana | DEC | MZ264241 | Burkholderiaceae | *Caballeronia* | 100 |
| INML1 | INML1 | A | M4 crypt | Indiana | MLF | MZ264270 | Burkholderiaceae | *Caballeronia* | 100 |
| INML2 | INML2 | A | M4 crypt | Indiana | MLF | MZ264242 | Burkholderiaceae | *Caballeronia* | 100 |
| INML3_ML3B | INML3 | A | M4 crypt | Indiana | MLF | MZ264243 | Burkholderiaceae | *Caballeronia* | 100 |
| INML3_ML3Y | INML3 | A | M4 crypt | Indiana | MLF | MZ264246 | Burkholderiaceae | *Caballeronia* | 100 |

| **Isolate Name** | **Individual** | **Life Stage** | **Tissue** | **State** | **Site** | **NCBI** | **Family** | ***Genus*** | **% Match to Genus** |
|---|---|---|---|---|---|---|---|---|---|
| INML5 | INML5 | A | M4 crypt | Indiana | MLF | MZ264244 | Burkholderiaceae | *Caballeronia* | 100 |
| INSB1 | INSB1 | A | M4 crypt | Indiana | SBG | MZ264245 | Burkholderiaceae | *Caballeronia* | 100 |
| NCF2_F2 | NCF2 | A | M4 crypt | North Carolina | FFA | MZ264248 | Burkholderiaceae | *Caballeronia* | 100 |
| NCF4 | NCF4 | A | M4 crypt | North Carolina | FFA | MZ264268 | Burkholderiaceae | *Caballeronia* | 100 |
| NCTM1 | NCTM1 | A | M4 crypt | North Carolina | TMF | MZ264247 | Burkholderiaceae | *Caballeronia* | 100 |
| NCTM5 | NCTM5 | A | M4 crypt | North Carolina | TMF | MZ264249 | Burkholderiaceae | *Caballeronia* | 100 |
| SB13_M4_a | SB13 | A | M4 crypt | Georgia | WOG | KT259185 | Burkholderiaceae | *Caballeronia* | 100 |
| SB14_M4_a | SB14 | A | M4 crypt | Georgia | WOG | KT259132 | Burkholderiaceae | *Caballeronia* | 100 |
| SB17_M4_b | SB17 | A | M4 crypt | Georgia | OAK | MH636870 | Burkholderiaceae | *Caballeronia* | 100 |
| SB18_M4_b | SB18 | A | M4 crypt | Georgia | OAK | KT259186 | Burkholderiaceae | *Caballeronia* | 100 |
| SB19_M4_a | SB19 | A | M4 crypt | Georgia | OAK | KT259133 | Burkholderiaceae | *Caballeronia* | 100 |
| SB19_M4_b | SB19 | A | M4 crypt | Georgia | OAK | KT259134 | Burkholderiaceae | *Caballeronia* | 100 |
| SB1b_LB | SB1 | A | M4 crypt | Georgia | CRY | KT259135 | Burkholderiaceae | *Caballeronia* | 100 |
| SB1B_V | SB1 | A | M4 crypt | Georgia | CRY | KT259136 | Burkholderiaceae | *Caballeronia* | 100 |
| SB1c_LB | SB1 | A | M4 crypt | Georgia | CRY | KT259137 | Burkholderiaceae | *Caballeronia* | 100 |
| SB1c_V | SB1 | A | M4 crypt | Georgia | CRY | KT259138 | Burkholderiaceae | *Caballeronia* | 100 |
| SB1d_LB | SB1 | A | M4 crypt | Georgia | CRY | KT259139 | Burkholderiaceae | *Caballeronia* | 100 |
| SB1e_LB | SB1 | A | M4 crypt | Georgia | CRY | KT259140 | Burkholderiaceae | *Caballeronia* | 100 |
| SB1e_V | SB1 | A | M4 crypt | Georgia | CRY | KT259141 | Burkholderiaceae | *Caballeronia* | 100 |
| SB1f_LB | SB1 | A | M4 crypt | Georgia | CRY | KT259142 | Burkholderiaceae | *Caballeronia* | 100 |
| SB1f_V | SB1 | A | M4 crypt | Georgia | CRY | KT259143 | Burkholderiaceae | *Caballeronia* | 100 |
| SB1g_LB | SB1 | A | M4 crypt | Georgia | CRY | KT259144 | Burkholderiaceae | *Caballeronia* | 100 |
| SB1g_V | SB1 | A | M4 crypt | Georgia | CRY | KT259145 | Burkholderiaceae | *Caballeronia* | 100 |
| SB1h_LB | SB1 | A | M4 crypt | Georgia | CRY | KT259146 | Burkholderiaceae | *Caballeronia* | 100 |
| SB22_M4_a | SB22 | A | M4 crypt | Georgia | CRY | KT259147 | Burkholderiaceae | *Caballeronia* | 100 |
| SB22_M4_b | SB22 | A | M4 crypt | Georgia | CRY | KT259187 | Burkholderiaceae | *Caballeronia* | 100 |

| **Isolate Name** | **Individual** | **Life Stage** | **Tissue** | **State** | **Site** | **NCBI** | **Family** | ***Genus*** | **% Match to Genus** |
|---|---|---|---|---|---|---|---|---|---|
| SB23_M4_c | SB23 | A | M4 crypt | Georgia | CRY | KT259183 | Burkholderiaceae | *Caballeronia* | 100 |
| SB23_M4_d | SB23 | A | M4 crypt | Georgia | CRY | KT259188 | Burkholderiaceae | *Caballeronia* | 100 |
| SB24_M4_a | SB24 | A | M4 crypt | Georgia | CRY | KT259148 | Burkholderiaceae | *Caballeronia* | 100 |
| SB25_M4_b | SB25 | A | M4 crypt | Georgia | CRY | KT259149 | Burkholderiaceae | *Caballeronia* | 100 |
| SB25_M4_c | SB25 | A | M4 crypt | Georgia | CRY | KT259150 | Burkholderiaceae | *Caballeronia* | 100 |
| SB26_M4_a | SB26 | A | M4 crypt | Georgia | CRY | KT259189 | Burkholderiaceae | *Caballeronia* | 100 |
| SB26_M4_b | SB26 | A | M4 crypt | Georgia | CRY | MH636872 | Burkholderiaceae | *Caballeronia* | 100 |
| SB27_M4_a | SB27 | A | M4 crypt | Georgia | CRY | KT259184 | Burkholderiaceae | *Caballeronia* | 100 |
| SB28_M4_a | SB28 | A | M4 crypt | Missouri | BCI | MH791154 | Burkholderiaceae | *Caballeronia* | 100 |
| SB29_M4_b | SB29 | A | M4 crypt | Missouri | BCI | KT259190 | Burkholderiaceae | *Caballeronia* | 100 |
| SB29_M4_c | SB29 | A | M4 crypt | Missouri | BCI | KT259151 | Burkholderiaceae | *Caballeronia* | 100 |
| SB30_M4_a | SB30 | A | M4 crypt | Missouri | BCI | KT259191 | Burkholderiaceae | *Caballeronia* | 100 |
| SB31_M4_b | SB31 | A | M4 crypt | Missouri | BCI | MH636871 | Burkholderiaceae | *Caballeronia* | 100 |
| SB7b | SB7 | A | M4 crypt | Georgia | NDG | KT259152 | Burkholderiaceae | *Caballeronia* | 100 |
| SB7d | SB7 | A | M4 crypt | Georgia | NDG | KT259153 | Burkholderiaceae | *Caballeronia* | 100 |
| SB7f | SB7 | A | M4 crypt | Georgia | NDG | KT259154 | Burkholderiaceae | *Caballeronia* | 100 |
| SB7g | SB7 | A | M4 crypt | Georgia | NDG | KT259155 | Burkholderiaceae | *Caballeronia* | 100 |
| SB7h | SB7 | A | M4 crypt | Georgia | NDG | KT259156 | Burkholderiaceae | *Caballeronia* | 100 |
| SB8b | SB8 | A | M4 crypt | Georgia | NDG | KT259157 | Burkholderiaceae | *Caballeronia* | 100 |
| SB8c | SB8 | A | M4 crypt | Georgia | NDG | KT259158 | Burkholderiaceae | *Caballeronia* | 100 |
| SB8d | SB8 | A | M4 crypt | Georgia | NDG | KT259159 | Burkholderiaceae | *Caballeronia* | 100 |
| SB8e | SB8 | A | M4 crypt | Georgia | NDG | KT259160 | Burkholderiaceae | *Caballeronia* | 100 |
| SB8f | SB8 | A | M4 crypt | Georgia | NDG | KT259161 | Burkholderiaceae | *Caballeronia* | 100 |
| SB8g | SB8 | A | M4 crypt | Georgia | NDG | KT259162 | Burkholderiaceae | *Caballeronia* | 100 |
| SQ4a | SQ4 | A | M4 crypt | Georgia | OAK | MH636869 | Burkholderiaceae | *Caballeronia* | 99 |
| SQ4f | SQ4 | A | M4 crypt | Georgia | OAK | KT259164 | Burkholderiaceae | *Caballeronia* | 93 |

| **Isolate Name** | **Individual** | **Life Stage** | **Tissue** | **State** | **Site** | **NCBI** | **Family** | ***Genus*** | **% Match to Genus** |
|---|---|---|---|---|---|---|---|---|---|
| SQ5a | SQ5 | A | M4 crypt | Georgia | OAK | KT259166 | Burkholderiaceae | *Caballeronia* | 100 |
| SQ5b | SQ5 | A | M4 crypt | Georgia | OAK | KT259167 | Burkholderiaceae | *Caballeronia* | 100 |
| SQ5c | SQ5 | A | M4 crypt | Georgia | OAK | KT259168 | Burkholderiaceae | *Caballeronia* | 100 |
| SQ5d | SQ5 | A | M4 crypt | Georgia | OAK | KT259169 | Burkholderiaceae | *Caballeronia* | 100 |
| SQ5e | SQ5 | A | M4 crypt | Georgia | OAK | KT259170 | Burkholderiaceae | *Caballeronia* | 100 |
| SQ5f | SQ5 | A | M4 crypt | Georgia | OAK | KT259171 | Burkholderiaceae | *Caballeronia* | 100 |
| SQ5g | SQ5 | A | M4 crypt | Georgia | OAK | KT259172 | Burkholderiaceae | *Caballeronia* | 100 |
| SQ5h | SQ5 | A | M4 crypt | Georgia | OAK | KT259173 | Burkholderiaceae | *Caballeronia* | 100 |
| SQ5i | SQ5 | A | M4 crypt | Georgia | OAK | KT259174 | Burkholderiaceae | *Caballeronia* | 100 |
| SQ6a | SQ6 | A | M4 crypt | Georgia | OAK | KT259175 | Burkholderiaceae | *Caballeronia* | 100 |
| SQ6b | SQ6 | A | M4 crypt | Georgia | OAK | KT259176 | Burkholderiaceae | *Caballeronia* | 100 |
| SQ6d | SQ6 | A | M4 crypt | Georgia | OAK | KT259177 | Burkholderiaceae | *Caballeronia* | 100 |
| SQ6e | SQ6 | A | M4 crypt | Georgia | OAK | KT259178 | Burkholderiaceae | *Caballeronia* | 100 |
| SQ6f | SQ6 | A | M4 crypt | Georgia | OAK | KT259179 | Burkholderiaceae | *Caballeronia* | 100 |
| SQ6g | SQ6 | A | M4 crypt | Georgia | OAK | KT259180 | Burkholderiaceae | *Caballeronia* | 100 |
| SQ6h | SQ6 | A | M4 crypt | Georgia | OAK | KT259181 | Burkholderiaceae | *Caballeronia* | 100 |
| SQ6i | SQ6 | A | M4 crypt | Georgia | OAK | KT259182 | Burkholderiaceae | *Caballeronia* | 100 |
| WG_1_5s_s | WG_1_5 | A | M4 crypt | Gerogia | WOG | MZ264236 | Burkholderiaceae | *Caballeronia* | 100 |

**Supplemental Table 2. Non-*Caballeronia* spp. bacteria isolated from *Anasa tristis.*** Abbreviations are as in Supplemental Table 1. Classification and % match to genus are based on the Ribosomal Database Project (RDP) classifier.

| Isolate Name | Individual | Location | Life Stage | Tissue | NCBI Accession | Phylum | Class | Order | Family | Genus | % Match to Genus |
|---|---|---|---|---|---|---|---|---|---|---|---|
| SB21_M4_a | SB21 | CRY | A | M4 crypt | MH828198 | Firmicutes | Bacilli | Bacillales | Bacillaceae 1 | *Bacillus* | 100 |
| SB23_M4_a | SB23 | CRY | A | M4 crypt | MH828199 | Firmicutes | Bacilli | Bacillales | Bacillaceae 1 | *Bacillus* | 100 |
| SB27_M4_b | SB27 | CRY | A | M4 crypt | MH828202 | Firmicutes | Bacilli | Bacillales | Bacillaceae 1 | *Bacillus* | 100 |
| SB28_M4_c | SB28 | BCI | A | M4 crypt | MH828205 | Firmicutes | Bacilli | Bacillales | Bacillaceae 1 | *Bacillus* | 100 |
| SB19_M4_c | SB19 | OAK | A | M4 crypt | MH828197 | Firmicutes | Bacilli | Bacillales | Paenibacillaceae 1 | *Paenibacillus* | 100 |
| SB27_M4_c | SB27 | CRY | A | M4 crypt | MH828203 | Firmicutes | Bacilli | Bacillales | Staphylococcaceae | *Staphylococcus* | 100 |
| SB23_M4_b | SB23 | CRY | A | M4 crypt | MH828200 | Firmicutes | Bacilli | Lactobacillales | Enterococcaceae | *Enterococcus* | 100 |
| A33_M4_g | A.33 | CRY | A | M4 crypt | MH828196 | Proteobacteria | Gammaproteobacteria | Enterobacteriales | Enterobacteriaceae | *Klebsiella* | 93 |
| SB31_M4_a | SB31 | BCI | A | M4 crypt | MH828207 | Proteobacteria | Gammaproteobacteria | Enterobacteriales | Enterobacteriaceae | *Klebsiella* | 94 |
| 2_61_a | 2.61 | CRY | 2 | Whole Body | MH828191 | Proteobacteria | Gammaproteobacteria | Enterobacteriales | Enterobacteriaceae | *Serratia* | 100 |
| 2_61_b | 2.61 | CRY | 2 | Whole Body | MH828192 | Proteobacteria | Gammaproteobacteria | Enterobacteriales | Enterobacteriaceae | *Serratia* | 100 |
| A33_M4_b | A.33 | CRY | A | M4 crypt | MH828193 | Proteobacteria | Gammaproteobacteria | Pseudomonadales | Moraxellaceae | *Acinetobacter* | 100 |
| SB28_M4_b | SB28 | BCI | A | M4 crypt | MH828204 | Proteobacteria | Gammaproteobacteria | Pseudomonadales | Pseudomonadaceae | *Pseudomonas* | 100 |
| A33_M4_e | A.33 | CRY | A | M4 crypt | MH828195 | Proteobacteria | Gammaproteobacteria | Xanthomonadales | Xanthomonadaceae | *Stenotrophomonas* | 100 |

**Supplemental Figure 1. 16S rRNA phylogeny of *Caballeronia* isolates from squash bug crypts, including symbionts of other true bugs species and other Burkholderiaceae.** Topology is the same as in Figure 3 in the main text. NCBI accession numbers for *A. tristis* symbionts are in Table S1. All other accession numbers follow taxa names. Shaded boxes indicate isolates used in fitness assays.

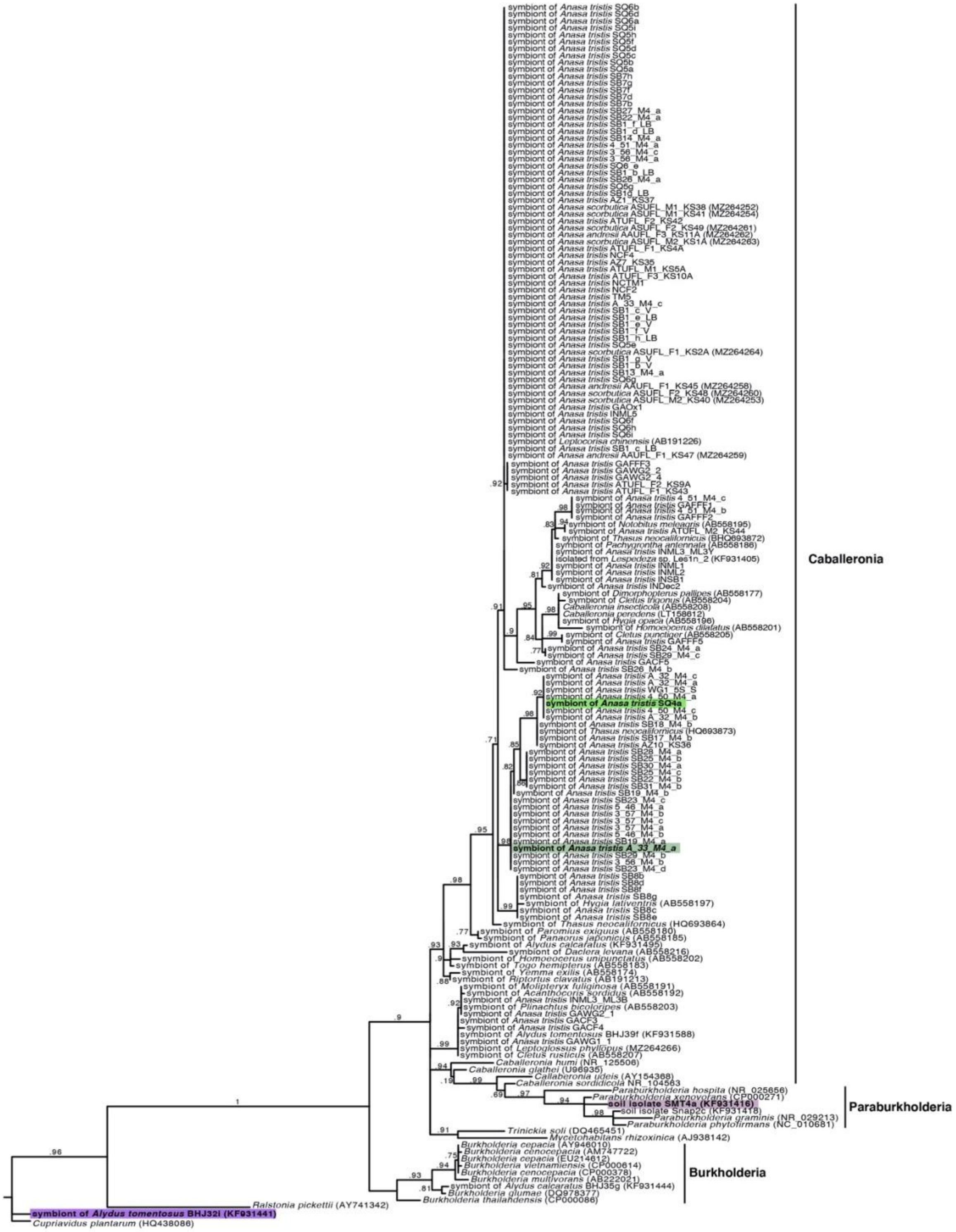

**Supplemental Figure 2. Set up for rearing squash bugs on plants.** Squash plants are grown from sterilized seeds in pots that extend into a plastic bin filled with water. The water is fertilized with Botanicare® Pure Blend Pro Grow Organic Fertilizer. The bin is filled with fertilized water until reaching the bottom of the pot.

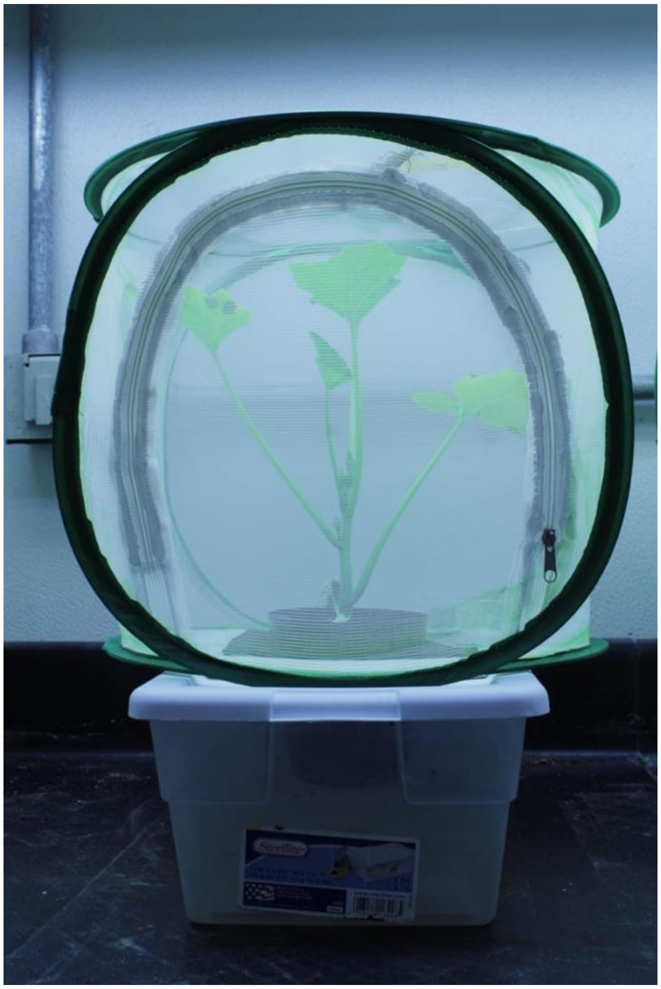

**Supplemental Table 3**. Isolation of GFP-labeled bacteria (SQ4a) from eggs washed in solution with alternative bacterial concentrations. CFUs indicates Colony Forming Units.

| **Estimated concentration of inoculation solution (CFUs per uL)** | **Total estimate of CFUs on sample of two eggs** | ***Caballeronia* detected?** |
|---|---|---|
| $5.8 \times 10^3$ | 97 | yes |
| $5.8 \times 10^3$ | 219 | yes |
| $5.8 \times 10^3$ | 580 | yes |
| $5.8 \times 10^2$ | 5 | yes |
| $5.8 \times 10^2$ | 14 | yes |
| $5.8 \times 10^2$ | 3 | yes |
| $3.5 \times 10^1$ | 1 | yes |
| $3.5 \times 10^1$ | 17 | yes |
| $3.5 \times 10^1$ | 2 | yes |
| 3.5 | 0 | no |
| 3.5 | 0 | no |
| 3.5 | 0 | no |

**Supplemental Figure 3. Estimated population size of *Caballeronia* SQ4a in adult female and male crypts.** Insects were reared on plants. Bars indicate means, and points indicate estimate for each individual.

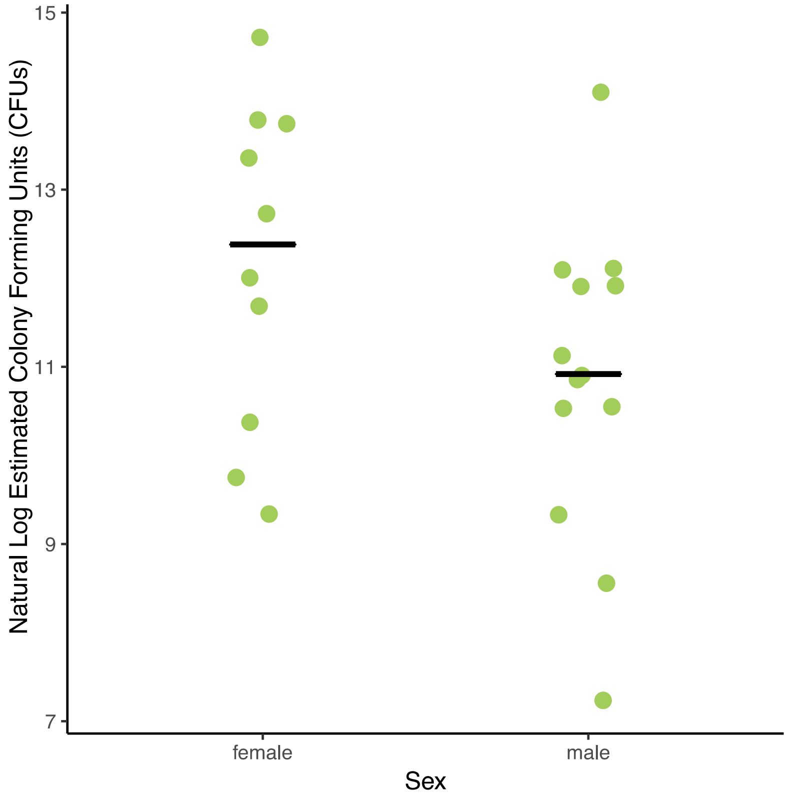

**Supplemental Table 4.** Results of Tukey's post-hoc comparisons of development time in days from hatch to each instar when fed alternative bacterial strains. Each cell contains the z-value. *indicates adjusted $P < 0.05$, ** adjusted $P < 0.01$, *** adjusted $P < 0.001$. All significant differences are highlighted in bold.

| Comparison | Hatch to 3rd Instar | Hatch to 4th Instar | Hatch to 5th Instar | Hatch to Adult |
|---|---|---|---|---|
| $H_2O$ - SQ4a | **3.18*** | **18.32***** | **27.01***** | **21.96***** |
| $H_2O$ - A33 | **3.36**** | **17.43***** | **25.05***** | **19.36***** |
| $H_2O$ - SMT4a | 0.13 | 0.50 | **6.61***** | **5.80***** |
| $H_2O$ - BHJ | 1.00 | -1.73 | 0.37 | -1.57 |
| SQ4a - A33 | 0.35 | 0.06 | -0.54 | -1.72 |
| SQ4a - SMT4a | **-2.88*** | **-17.70***** | **-19.06***** | **-14.76***** |
| SQ4a - BHJ | -2.44 | **-19.90***** | **-26.30***** | **-23.25***** |
| A33 - SMT4a | **-3.07*** | **-16.94***** | **-17.68***** | **-12.64***** |
| A33 - BHJ | -2.67 | **-18.92***** | **-24.42***** | **-20.65***** |
| SMT4a - BHJ | 0.57 | -1.10 | **-6.17***** | **-7.19***** |

**Supplemental Table 5.** Results of Tukey's post-hoc comparisons of impact of bacterial inoculation treatment on adult pronotal width. Each cell contains the z-value. *indicates adjusted $P < 0.05$, ** adjusted $P < 0.01$, *** adjusted $P < 0.001$. All significant differences are highlighted in bold.

| Comparison | Difference in Means (log-transformed pronotal width, mm) |
|---|---|
| $H_2O$ – SQ4a | **-0.06***** |
| $H_2O$ - A33 | **-0.06***** |
| $H_2O$ – SMT4a | -0.02 |
| $H_2O$ – BHJ | -0.01 |
| SQ4a - A33 | 0.002 |
| SQ4a – SMT4a | **0.04***** |
| SQ4a – BHJ | **0.05***** |
| A33 – SMT4a | **0.04***** |
| A33 – BHJ | **0.05***** |
| SMT4a – BHJ | 0.01 |
